# Heterogeneity of monocyte subsets and susceptibility to influenza virus contribute to inter-population variability of protective immunity

**DOI:** 10.1101/2020.12.07.414151

**Authors:** Mary O’Neill, Hélène Quach, Julien Pothlichet, Yann Aquino, Aurélie Bisiaux, Nora Zidane, Matthieu Deschamps, Valentina Libri, Milena Hasan, Shen-Ying Zhang, Qian Zhang, Daniela Matuozzo, Aurélie Cobat, Laurent Abel, Jean-Laurent Casanova, Nadia Naffakh, Maxime Rotival, Lluis Quintana-Murci

**Affiliations:** Human Evolutionary Genetics Unit, Institut Pasteur, UMR 2000, CNRS, Paris 75015, France; Muséum National d’Histoire Naturelle, UMR7206, CNRS, Université de Paris, Paris 75016, France; DIACCURATE, Paris, 75014, France; Biodiversity and Epidemiology of Bacterial Pathogens Unit, Institut Pasteur, Paris 75015, France; Cytometry and Biomarkers UTechS, Institut Pasteur, Paris 75015, France; St. Giles Laboratory of Human Genetics of Infectious Diseases, Rockefeller Branch, The Rockefeller University, New York, NY, 10065, USA; Laboratory of Human Genetics of Infectious Diseases, Necker Branch, Necker Hospital for Sick Children, INSERM UMR 1163, Necker Hospital for Sick Children, Paris 75015, France; Paris University, Imagine Institute, Paris 75015, France; Howard Hughes Medical Institute, New York, NY, USA; RNA Biology of Influenza Virus Unit, Institut Pasteur, Paris, 75015, France; Chair of Human Genomics and Evolution, Collège de France, Paris, 75005, France

**Author notes:** Correspondence and requests for materials should be addressed to M.R or L.Q.-M.,). M.R. and L.Q.M. contributed equally to this work. **Authors contributions:** M.B.O, M.R. and L.Q.M. conceived and designed the study. M.B.O., H.Q., J.P., Y.A., A.B., N.Z., and M.D. conducted the experiments at the Human Evolutionary Genetics Unit. M.B.O., M.R., Y.A. and D.M. developed computational methods and performed bioinformatic analyses. J.P., V.L., M.H., S.Z.Y., Q.Z., A.C. L.A., J.L.C. and N.N. provided resources, expertise and feedback. M.R. and L.Q.M supervised the study. L.Q.M secured funding. M.B.O., M.R. and L.Q.M. wrote the manuscript with input from all authors. **Competing Interest Statement:** The authors declare no competing interests.

**Keywords:** Influenza, monocytes, single-cell, transcriptomics, ancestry

## Abstract

There is considerable inter-individual and inter-population variability in response to viruses. The potential of monocytes to elicit type-I interferon responses has attracted attention to their role in viral infections. Here, we use an *ex vivo* model to characterize the role of cellular heterogeneity in human variation of monocyte responses to influenza A virus (IAV) exposure. Using single-cell RNA-sequencing, we show widespread inter-individual variability in the percentage of IAV-infected monocytes. We show that cells escaping viral infection display increased mRNA expression of type-I interferon stimulated genes and decreased expression of ribosomal genes, relative to both infected cells and those never exposed to IAV. While this host defense strategy is shared between *CD16*^*+*^/*CD16*^*-*^ monocytes, we also uncover *CD16*^*+*^-specific mRNA expression of *IL6* and *TNF* in response to IAV, and a stronger resistance of *CD16*^*+*^ monocytes to IAV infection. Notably, individuals with high cellular susceptibility to IAV are characterized by a lower activation at basal state of an IRF/STAT-induced transcriptional network, which includes antiviral genes such as *IFITM3, MX1*, and *OAS3*. Finally, using flow cytometry and bulk RNA-sequencing across 200 individuals of African and European ancestry, we observe a higher number of *CD16*^+^ monocytes and lower susceptibility to IAV infection among monocytes from individuals of African-descent. Collectively, our results reveal the effects of IAV infection on the transcriptional landscape of human monocytes and highlight previously unappreciated differences in cellular susceptibility to IAV infection between individuals of African and European ancestry, which may account for the greater susceptibility of Africans to severe influenza.

**Significance Statement:** Monocytes may play a critical role during severe viral infections. Our study tackles how heterogeneity in monocyte subsets and activation contributes to shape individual differences in the transcriptional response to viral infections. Using single-cell RNA-sequencing, we reveal heterogeneity in monocyte susceptibility to IAV infection, both between *CD16*^*+*^/*CD16*^*-*^ monocytes and across individuals, driven by differences in basal activation of an IRF/STAT-induced antiviral program. Furthermore, we show a decreased ability of IAV to infect and replicate in monocytes from African-ancestry individuals, with possible implications for antigen presentation and lymphocyte activation. These results highlight the importance of early cellular activation in determining an individuals’ innate immune response to viral infection.

## Introduction

Respiratory viruses with pandemic potential pose enormous health and economic impacts on the human population. In the last century, we have witnessed outbreaks of several coronaviruses, including SARS-CoV-2, SARS-CoV-1 and MERS, and a number of avian and swine influenza A viruses (IAV). A particularly harrowing and shared feature of these pandemics are the sudden deaths of otherwise healthy individuals (1). A hyperinflammatory state characterized by high levels of inflammatory cytokines, often referred to as a ‘cytokine storm’ (2, 3), has emerged as a hallmark of these severe viral infections. While still controversial, there is increasing evidence to suggest that the mononuclear phagocyte system is an important immunological determinant of this phenotype (4-6). Upon viral infection, sentinel cells such as lung-resident macrophages trigger complex signaling cascades that recruit leukocytes to the site of infection, among them monocytes. These infiltrating monocytes differentiate into monocyte-derived dendritic cells or macrophages, enabling viral clearance through the induction of the adaptive response, and help replenish the pool of tissue-resident alveolar macrophages (4, 7).

In humans, circulating monocytes are divided into classical (∼80%), intermediate (∼15%), and nonclassical (∼5%) subsets, based on surface receptor expression of the cluster-determinant antigens CD14 and CD16 (8). While nonclassical monocytes (CD14^+^CD16^++^) are long-lived and ‘patrol’ healthy tissues through long-range crawling on the endothelium, classical (CD14^++^CD16^-^) and intermediate (CD14^++^CD16^+^) monocytes are recruited to the lung in response to viral infection, where they secrete inflammatory cytokines and chemokines, as well as type I interferons (IFNs) (7, 9-11). In most individuals, recruited cells help clear infection despite being susceptible to infection themselves (12, 13); yet, in some individuals, a dysfunctional immune response occurs resulting in widespread lung inflammation. Whether monocyte subsets behave differently upon viral exposure, and how direct viral sensing and exposure to secreted cytokines shape monocyte activation and differentiation are not well understood.

Variation in blood composition and cellular proportions have been shown to be one of the main factors underlying transcriptional variation in immune genes across individuals (14), with these proportions being influenced by both genetic and non-heritable factors (15-17). Recently, we characterized the genetic architecture of transcriptional responses of primary monocytes from 200 individuals of African and European ancestry to *ex vivo* challenge with viral stimuli (18). In this model, where we were able to control for viral determinants of disease (i.e. dose and strain), we reported marked inter- and intra-population differences in transcriptional responses to IAV. While our analyses revealed numerous *cis*-expression quantitative trait loci (18), genetic variants could only account for a small fraction of expression variation, in line with other studies (14, 19).

Here, we implemented single-cell RNA-sequencing (scRNA-seq) on human primary monocytes exposed to IAV to investigate (i) the effects of direct viral infection versus activation by exposure to secreted cytokines, (ii) the subset-specific responses of monocytes to viral challenge, and (iii) the extent of interindividual and between-population variation in the proportions of monocyte subsets and the degree of monocyte susceptibility to IAV infection.

## Results

### Using single-cell RNA-sequencing to investigate cellular heterogeneity

To investigate the role of cellular heterogeneity in driving immune variability across individuals, we performed a time-course experiment where we monitored the CD14^+^ fraction of peripheral blood mononuclear cells (PBMCs) from eight donors, both in the presence and absence of viral challenge. To maximize inter-individual variability, we chose individuals from two distinct ancestries whose cells demonstrated extreme responses to viral stimuli in a previous bulk RNA-seq experiment (18). Droplet-based scRNA-seq was performed on monocytes from all eight donors immediately before infection initiation (T_0_), as well as at 2 (T_2_), 4 (T_4_), 6 (T_6_), and 8 (T_8_) hours post challenge with A/USSR/90/1977(H1N1) at a multiplicity of infection (MOI) equal to 1 (IAV-challenged) and mock infection (non-infected). To mitigate batch effects, we pooled IAV-challenged and non-infected cells from distinct donors in each library, assigning cells to their condition in silico via genetic barcoding (20). After stringent quality control, our final dataset contained 88,559 high-quality cells, among which we predicted >99% monocyte purity at T_0_ (**Fig. 1*A*; *SI Appendix*, Figs. S1 and S2**). At later time points, a substantial fraction of non-infected cells (up to 70% at T_8_) were predicted to be macrophage-like, indicating monocyte differentiation over the course of the experiment. For clarity, we refer to cells as monocytes at T_0_ and as monocyte-derived cells from T_2_-T_8_.

**Fig. 1.**
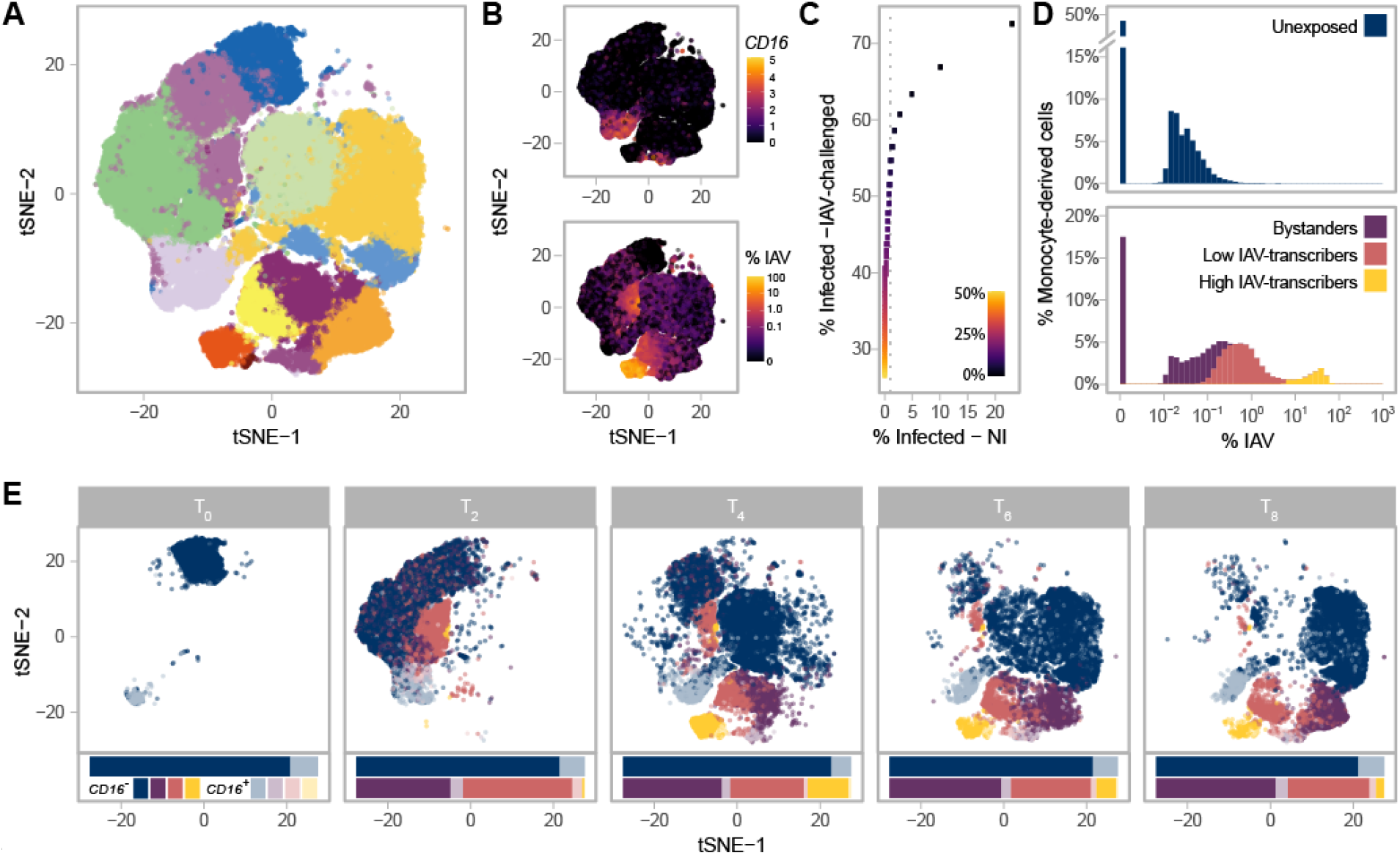
Single-Cell RNA-Sequencing of 88,559 Monocytes and Their Derived Cells. (***A***) Post-QC tSNE colored by unsupervised graph-based clusters. (***B***) Post-QC tSNE colored by *FCGR3A* (*CD16*) log_2_ normalized counts (top), or percentage of viral mRNAs (bottom). (***C***) Determination of the maximum contamination fraction by ambient RNA. The number of non-infected cells deemed to significantly express IAV transcripts (presumed false positives) versus the number of IAV-challenged cells deemed to significantly express IAV transcripts across a range of maximum contamination fractions from 1-50% (color bar). Dotted grey line is drawn at 1% on the x-axis. A maximum contamination fraction of 10% results in 1% of non-infected cells being classified as infected (false positive proxy), and half of IAV-challenged cells showing evidence of viral transcription. (***D***) Distribution of counts of viral origin across all donors, from T_2_ to T_8_. Cells are shown separately for non-infected (top) and IAV-challenged (bottom) conditions. Fill color reflects the cell state assignments. Note that the threshold used to define infected cells is dependent on the number of viral mRNAs in the ambient pool, and varies across libraries. (***E***) Post-QC tSNE stratified by time point. For each time point, cells are colored according to their *CD16*^*+/-*^ status (see key) and their assigned cell state (same as depicted in *D*). For each condition and time point, stacked bar charts below the tSNE represent the relative proportions of the various cell states and subsets.

### Stable *FCGR3A* expression distinguishes monocyte subsets over time

We next sought to characterize each cell by its mRNA expression of the canonical monocyte markers, CD14 and CD16, given that much of the structure in our data was associated with *FCGR3A* (aka *CD16*) mRNA expression. In droplet-based scRNA-seq, encapsulation of ambient mRNAs emanating from dying cells can occur during library preparation leading to spurious mRNA detection (21). We thus used a statistical framework to test whether *CD14* and *CD16* were expressed at a level significantly higher than expected when accounting for potential contamination from the ambient pool (***Methods***). Despite having been positively selected for the CD14 antigen, only 32.4% of monocytes significantly expressed *CD14* at T_0_; this percentage further decreased at later time points and remained <15% across all time points and conditions (average 6.4% s.d.: 5.0%, ***SI Appendix*, Fig. S3*A*-*C***). On the other hand, 12.1% of monocytes significantly expressed *FCGR3A* (*CD16*) (referred to as *CD16*^*+*^) at T_0_, this marker proving much more stable across conditions and time points (9.3% of *CD16*^*+*^ cells on average, s.d.: 1.8%, **Fig. 1*B* and *SI Appendix*, Fig. S3*D*-*F***). While we deciphered classical, intermediate, and nonclassical monocytes subsets at T_0_ (***SI Appendix*, Note 1 and Fig. S4; Dataset S1**), we focus on the simpler distinction of *CD16*^*-*^ and *CD16*^*+*^ subsets given that positive-selection for monocytes does not capture the entire nonclassical population and that we were unable to distinguish the intermediate and noncanonical subsets after T_0_.

### Functional features of monocyte subsets are conserved upon manipulation

To assess how transcriptional profiles of *CD16*^*-*^ and *CD16*^*+*^ monocytes and their derived-cells differ, we focused on the 5,681 genes expressed with a normalized log count > 0.1 in at least one condition, time point, and subset (**Dataset S2*A***). We found that the log_2_ fold change in gene expression between *CD16*^*+/-*^ subsets remained relatively stable over the course of the experiment (Pearson r between time points >0.42, and >0.52 for the non-infected and IAV-challenged conditions respectively, *p*-values < 2.2×10^−16^; ***SI Appendix*, Fig. S5*A***), and differentially expressed genes between *CD16*^*+/-*^ subsets were largely the same across conditions (Pearson r = 0.92, *p*-value < 2.2×10^−16^; ***SI Appendix*, Fig. S5*B***). We thus searched for genes that were consistently differentially expressed between *CD16*^*+*^ and *CD16*^*-*^ cells across all time points (including T_0_), conditions, and donors. We identified 266 genes over-expressed (log_2_FC>0.2, FDR<1%) in *CD16*^*+*^ cells relative to *CD16*^*-*^ cells, and 389 genes that showed the opposite pattern, and performed a GO-term enrichment analysis on these genes (**Dataset S2*B***). Consistent with previous reports (22-24), *CD16*^*-*^ subsets were characterized by high expression of several proinflammatory *S100 Calcium Binding Proteins* (*S100A12, S100A9*, and *S100A8*), contributing to a sizable GO-term enrichment in the defense response to fungus pathway (GO:0050832: OR=41.3, FDR=4.9×10^−4^), while *CD16*^*+*^ subsets were characterized by high expression of Fc-gamma receptor signaling pathway genes (GO:0038096: OR=8.7, FDR=6.2×10^−6^). Notably, *CD16*^*+*^ subsets over-expressed several type I IFN stimulated genes (ISGs) relative to *CD16*^*-*^ subsets (e.g. GO:000071357: OR=5.3, FDR=2.6×10^−3^), including the well-known viral restriction factors *IFITM3* and *OAS1*. Collectively, these results demonstrate *CD16* is a reliable marker at the mRNA level and that *CD16*^*+/-*^ monocyte subsets maintain functional differences upon manipulation.

### scRNA-seq highlights heterogeneity in monocyte susceptibility and viral transcription

Using the presence of IAV transcripts as a proxy for infection (**Fig. 1*B***), we next sought to distinguish cells that were successfully infected from those that were not. Among monocyte-derived cells that were exposed to IAV, we found that 50.3% expressed IAV transcripts above ambient levels when allowing up to 10% of mRNAs to come from the ambient pool. In contrast, less than 1% of non-infected cells showed evidence of viral transcription, supporting the validity of the threshold used to detect IAV expressing cells (**Fig. 1*C***). We deemed cells with statistical evidence for expression of IAV transcripts from the IAV-challenged condition as ‘infected’, while the remaining cells from this condition were considered as ‘bystanders’, as these either did not come into contact with the virus or were able to fully repress viral mRNA transcription. When comparing the percentage of infected cells between subsets, we noticed that *CD16*^*+*^ cells were slightly less likely to be infected than *CD16*^*-*^ cells (42.3% sd: 4.0% for *CD16*^*+*^ relative to 49.4% sd: 5.4% for *CD16*^*-*^, generalized linear model with CD16^+/-^ status, donor, and time point as covariates, *p*-value=0.006). This suggests a higher resistance of *CD16*^*+*^ to IAV infection, possibly related to the higher expression of ISGs observed in this subset (**Dataset S2 *A* and *B***).

We observed that the proportions of viral mRNAs among infected cells were bimodally distributed and largely varied between the clusters identified in our unsupervised analysis (**Fig. 1*D***). We used a Gaussian mixture model to locate the two modes of the distribution and further sub-classify infected cells into those with lower IAV mRNA levels (<1-6%) and those with higher IAV mRNA levels (6-83%); while viral mRNA levels are dictated by both the rate of transcription and degradation, for simplicity we refer to these infected cell states as ‘low IAV-transcribers’ and ‘high IAV-transcribers’, respectively. The proportions of infected cells among individuals remained largely unchanged over the course of the experiment; however, high-IAV transcribers were virtually absent at 2h (<2% of infected cells), peaked to ∼36% of IAV-infected cells at 4h, and decreased to 8.5% by 8h, suggesting that high-IAV transcribers represent a transient state of IAV-infection preceding IAV-induced apoptosis (**Fig. 1*E***). These results reveal profound heterogeneity in monocyte susceptibility and subsequent viral transcription upon IAV-challenge.

### Interplay of cytokine and ribosome networks drive cell states upon infection

To characterize host transcriptional responses over time, we next subsampled each subset (*CD16*^-^/*CD16*^+^), cell state (unexposed, bystander, infected), and time point in our scRNA-seq data to 100 cells, while ensuring a balanced representation of all donors. We then focused on the 6,669 host genes with average log_2_ normalized count >0.1 in at least one subgroup (**Dataset S3*A***). Overall, *CD16*^-^ and *CD16*^+^ subsets behaved similarly upon stimulation with changes in gene expression between cell states being strongly correlated among subsets (Pearson r = 0.83-0.95, *p*-values < 2.2×10^−16^; ***SI Appendix*, Fig. S6)**. GO term enrichment analyses of shared responses (FDR<1% & log_2_FC >0.2 in same direction in both subsets) uncovered several functional categories interacting to shape the activation state of cells (**Fig. 2*A***; **Dataset S3*B***). Both bystander and infected cells showed increased mRNA expression of genes involved in antigen processing and presentation via class I MHC (GO:0019885, OR=53.7, FDR < 2.0×10^−6^) and response to type I IFN (GO:0034340, OR >14.8, FDR < 3.3×10^−20^). Yet, bystander cells showed increased mRNA expression of type I IFN response and defense response to virus pathways relative to infected cells (GO:0034340: OR=13.4, FDR=4.4×10^−7^; GO:0051607: OR=9.0, FDR=2.1×10^−7^), while infected cells displayed higher mRNA expression of mitochondrial (GO:0005743, OR=4.7, FDR=3.3×10^−3^) and ribosomal genes (GO: 0005840, OR=117, FDR=1.0×10^−78^).

**Fig. 2.**
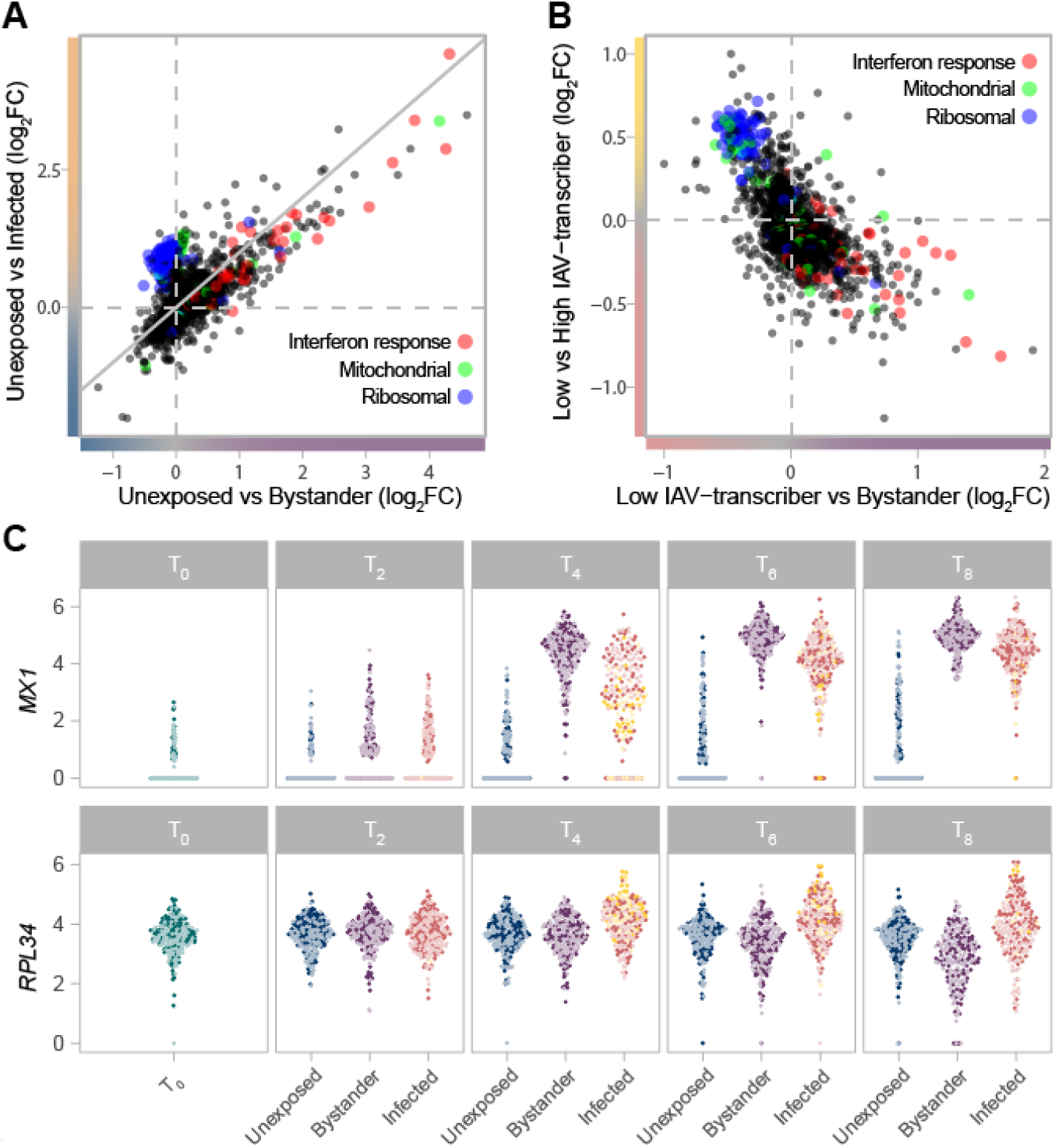
Gradient of mRNA Expression from Ribosomal and IFN-Stimulated Genes Separates Bystander and Infected Cells. (***A***) Transcriptional responses of cells upon IAV-challenge (T_2_-T_8_) highlight the interplay between IFN response (GO:0034340), ribosomal (GO:0005840), and mitochondrial (GO:0005743) genes. The log_2_FC change in gene expression between unexposed and bystander cells is plotted on the x-axis, while the log_2_FC change in gene expression between unexposed and infected cells is plotted on the y-axis. Values are plotted based on a meta-analysis across time points and subsets, based on a subsampled data set with balanced representation of all donors. (***B***) The interplay between IFN response (GO:0034340), ribosomal (GO:0005840), and mitochondrial (GO:0005743) genes among cells exposed to IAV. The log_2_FC change in gene expression between low IAV-transcribing infected and bystander cells is plotted on the x-axis, while the log_2_FC change in gene expression between low IAV-transcribing infected and high IAV-transcribing infected cells is plotted on the y-axis. Values are plotted based on a meta-analysis across monocyte subsets at T_4_. (***C***) mRNA expression levels of representative IFN-stimulated (*MX1*) and ribosomal (*RPL34*) genes across the subsampled dataset. Colors reflect the cell state and subset assignment depicted in **Fig. 1 *D*** and ***E***.

Among infected cells, ribosomal genes showed higher activity among high IAV-transcribing cells relative to low IAV-transcribing cells (**Fig. 2*B***, comparison only made at T_4_ due to sample size constraints, e.g. GO:0019083: OR=137, FDR=6.1×10^−65^). This observation is consistent with the notion that the expression of viral proteins is dependent on cellular ribosomes, with recent data suggesting that IAVs do not induce a global shut-off of cellular translation but rather a reshaping of the translation landscape (25-27). Likewise, among bystander cells, numerous ribosomal genes were downregulated at later time points relative to unexposed cells (**Fig. 2*A* and *C***; GO:0019083, OR=5.3, FDR=4.2×10^−6^), suggesting that repression of ribosomal subunits plays an active role in limiting viral replication. Collectively, these results suggest that ISGs and ribosomal expression interact to shape cell states upon IAV-challenge.

### Increased IRF and STAT activity drives stronger antiviral response

Despite qualitatively similar responses to infection between CD16^-^/CD16^+^ subsets (***SI Appendix*, Fig. S6**), we hypothesized that subtle differences in the intensity of such responses might contribute to the increased resistance of *CD16*^*+*^ cells to infection. We thus performed an interaction test on the subsampled scRNA-seq data, and searched for genes for which transcriptional response upon IAV-challenge differed between *CD16*^*-*^ and *CD16*^*+*^ subsets in either infected and/or bystander cells (***SI Appendix*, Fig. S6 *A* and *B***; **Dataset S3*A***). At FDR≤1%, we identified a total of 335 such genes, of which 98 differed between subsets only in bystander cells, 144 only in infected cells, and 93 in both. Hierarchical clustering highlighted eight major patterns of transcriptional responses (modules) among the 335 genes, several of which were associated with specific biological functions (**Fig. 3*A*; Dataset S3 *C* and *D***). Notably, module 1 (green) was enriched for genes in the antiviral response pathway (GO:0051607, OR=23.2, FDR<5.43×10^−7^) and displayed a stronger response in infected *CD16*^*+*^ cells relative to *CD16*^*-*^ infected cells. Of additional interest was the transient *CD16*^*+*^ specific transcription of the inflammatory cytokine genes *IL6* and *TNF*, following viral challenge (**Fig. 3*B***). We also found that several genes involved in the regulation and production of IL-6 and TNFα are over-expressed in *CD16*^*+*^ subsets at all time-points and conditions (**Dataset S2*B***), but only see active transcription of the cytokines upon viral exposure. These results reveal the strong antiviral and inflammatory potential of *CD16*^*+*^ relative to *CD16*^*-*^ monocytes in response to viral infection (28).

**Fig. 3.**
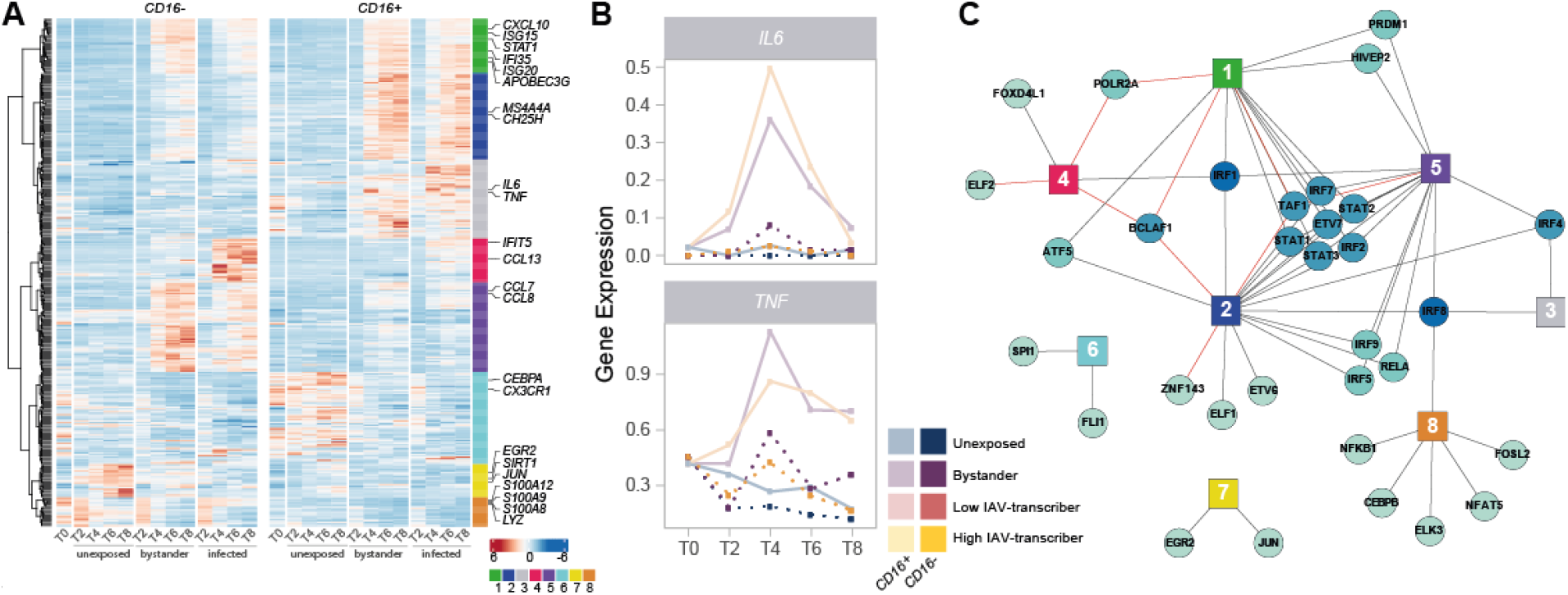
IRFs and STATs Have a Central Role in the Subset-Specific Responses to IAV Infection. (***A***) Heatmap of scaled gene expression from 335 genes displaying a subset-specific response to infection challenge. Genes are grouped into 8 modules based on hierarchical clustering of their expression patterns. Representative genes from each module are labelled. (***B***) Mean expression over time of *IL6* and *TNF*, across the different monocyte subsets and cell states. (***C***) Network of transcription factors (round nodes) associated with each gene expression module (square nodes). Transcription factor nodes are colored according to the number of modules they are associated with. Black lines represent enrichments of the module in TF targets, while red lines represent depletions.

We next sought to characterize the regulatory architecture underlying the 335 genes whose transcriptional response to IAV-challenge differed between monocyte subsets. Using SCENIC (29), we identified 113 high-confidence gene regulatory networks, or ‘regulons’, which are active in non-infected and/or IAV-challenged cells, each composed of a transcription factor (TF) and a set of predicted targets (genes). We used these 113 regulons to search for an enrichment/depletion of TF targets among the eight modules of genes displaying subset-specific response to infection (**Dataset S3*E***). Among modules associated with an increased expression in cells exposed to IAV (modules 1-5), we observed a widespread over-representation of targets of IFN regulatory factors (IRFs) and signal transducing and activators of transcription (STATs) (**Fig. 3*C***), reinforcing the central role of the IFN response upon IAV challenge. Interestingly, several of these factors displayed subset-specific activity themselves in response to IAV (*IRF1/2/7* and *STAT1/2/3*, FDR<1%), mirroring the expression patterns of module 1 (Pearson r>0.92). These results collectively highlight a *CD16*^*+*^-specific inflammatory response upon IAV-challenge and suggest stronger activation of IRF and STAT transcription factors as driver of the increased antiviral response observed in *CD16*^*+*^ cells upon IAV infection.

### Basal activation differences correlate with monocyte susceptibility

To explore the degree of inter-individual variation upon viral challenge, we next quantified IAV transcripts in the monocyte-derived cells of each individual, and created pseudo-bulk estimates by averaging the percent of viral mRNAs per-cell across all cells from each donor at each time point (**Fig. 4*A***). While viral mRNAs peaked at the same time for all individuals, we observed extensive variation in the levels of viral mRNAs and percentages of infected cells across individuals (**Fig. 4*B***). To identify specific genes that might underlie infection potential, we focused on the 4,589 genes that were expressed at > 0.1 log_2_ normalized counts in at least one canonical monocyte subset at T_0_. We identified a total of 3,131 genes that differed among our eight donors in either classical, intermediate, and/or nonclassical monocyte subsets (Kruskal-Wallis Rank Test, FDR 1%; **Dataset S4*A***). Within each subset, focusing on genes that significantly differ between donors, we searched for those for which mean expression at basal state was correlated with the percentage of infected cells at T_4_ among our eight donors. Despite our limited sample size, we found that cellular susceptibility was strongly correlated with basal expression of the well-known host viral restriction factor *IFITM3*. Although it reached significance only in nonclassical monocytes (FDR∼1%), the association remained strong in other subsets (*p* < 4.1 ×10^−4^; **Fig. 4*C***).

**Fig. 4.**
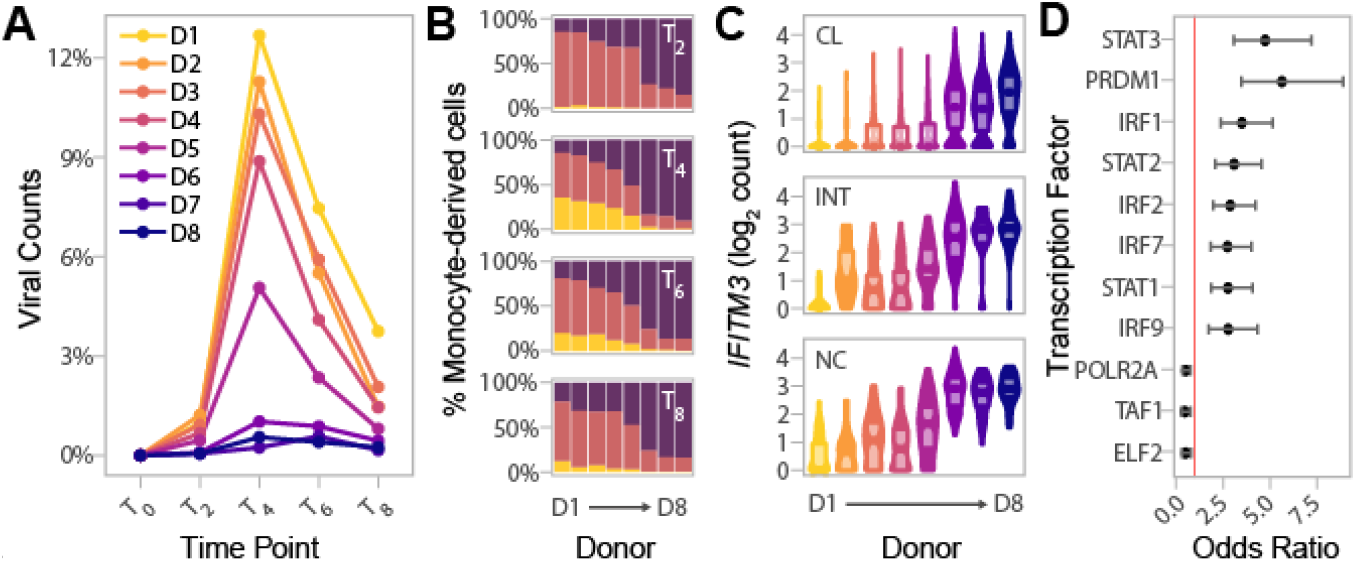
Basal IRF/STAT-Induced Transcriptional Network Underlies Inter-Individual Differences in Monocyte Susceptibility and IAV levels. (***A***) Pseudo-bulk estimates of the percentage of counts of viral origin in IAV-challenged condition (T_2_-T_8_). Donors are colored based on the rank of these pseudo-bulk estimates at the peak of viral transcription, T_4_, from that with highest observed viral mRNA level (D1) to that of the lowest (D8). (***B***) Proportions of cell states from the IAV-challenged condition at T_2_, T_4_, T_6_, and T_8_, in the eight donors. X-axis is ordered by decreasing viral mRNA levels found at T_4_ (D1-D8). (***C***) Log normalized expression values of *IFITM3* across all cells, stratified by canonical monocyte subsets, and separated by donor. Colors reflect the different donors depicted in ***A***. For each donor and monocyte subset, the violin plots show the full distribution of *IFITM3* expression across individual cells and boxplots highlight the median and interquartile range. (***D***) Enrichment of SCENIC-predicted targets among the 118 genes whose basal expression at T_0_ correlates with the percentage of infected cells at later time points (odds ratio and 95% confidence interval). Red line designates an odds ratio equal to 1. Only TFs significantly enriched among the 118 candidate genes are shown (FDR < 0.05). Abbreviations: classical (CL), intermediate (INT), and nonclassical (NC).

We next relaxed our search to all genes for which basal expression showed nominal correlation (*p* < 0.01) with the percentage of infected cells at T_4_. Depending on the monocyte subset, between 3.6 to 8.3% of genes matched these criteria, resulting in a set of 118 genes displaying correlation with monocyte susceptibility in at least one subset. These 118 genes were collectively enriched for several related biological processes such as defense response to virus (GO:0051607, OR=15.3, FDR=9.2×10^−19^) and response to type I IFN (GO:0034340, OR=19.6 FDR=8.4×10^−15^) (**Dataset S4*B***). Among genes contributing to this enrichment, we found additional antiviral genes such as *OAS3*, and *MX1*, as well as the critical TF, *IRF7*, involved in the severity of IAV-infection both in mice and humans (30-32). Finally, overlap with the TF targets identified by SCENIC revealed strong enrichments of several IRFs and STATs among the 118 genes, including IRF7, as well as STAT1, STAT2 and IRF9 that form the tripartite IFN-stimulated gene factor 3 (ISGF3) (**Fig. 4*D***; **Dataset S4*C***). Together, our results provide evidence that the basal mRNA expression of genes related to IFN-induced and antiviral responses are indicative of the proportion of cells that will become infected in the first cycle of IAV infection.

### African-ancestry monocytes are more resistant to infection

Lastly, we wondered how our findings of inter-individual variation might extrapolate to the population level. In a previous study (18), we challenged the primary monocytes from 200 Belgian individuals of African (AFB) and European (EUB) ancestry with the same IAV strain and MOI used in the present study, and performed bulk RNA-seq at 6 hours post infection (hpi). While basal (T_0_) expression profiles were not collected, flow cytometry labelling of CD14 and CD16 was performed on the CD14^+^-selected monocytes for the majority of donors. Interestingly, AFB individuals had higher proportions of *CD16*^*+*^ cells than EUB individuals (**Fig. 5*A***; ***SI Appendix*, Fig. S7**). In light of our findings that *CD16*^*+*^ cells are more resistant to IAV infection, we hypothesized that this might translate to lower infection rates among AFB monocytes relative to EUB monocytes.

**Fig. 5.**
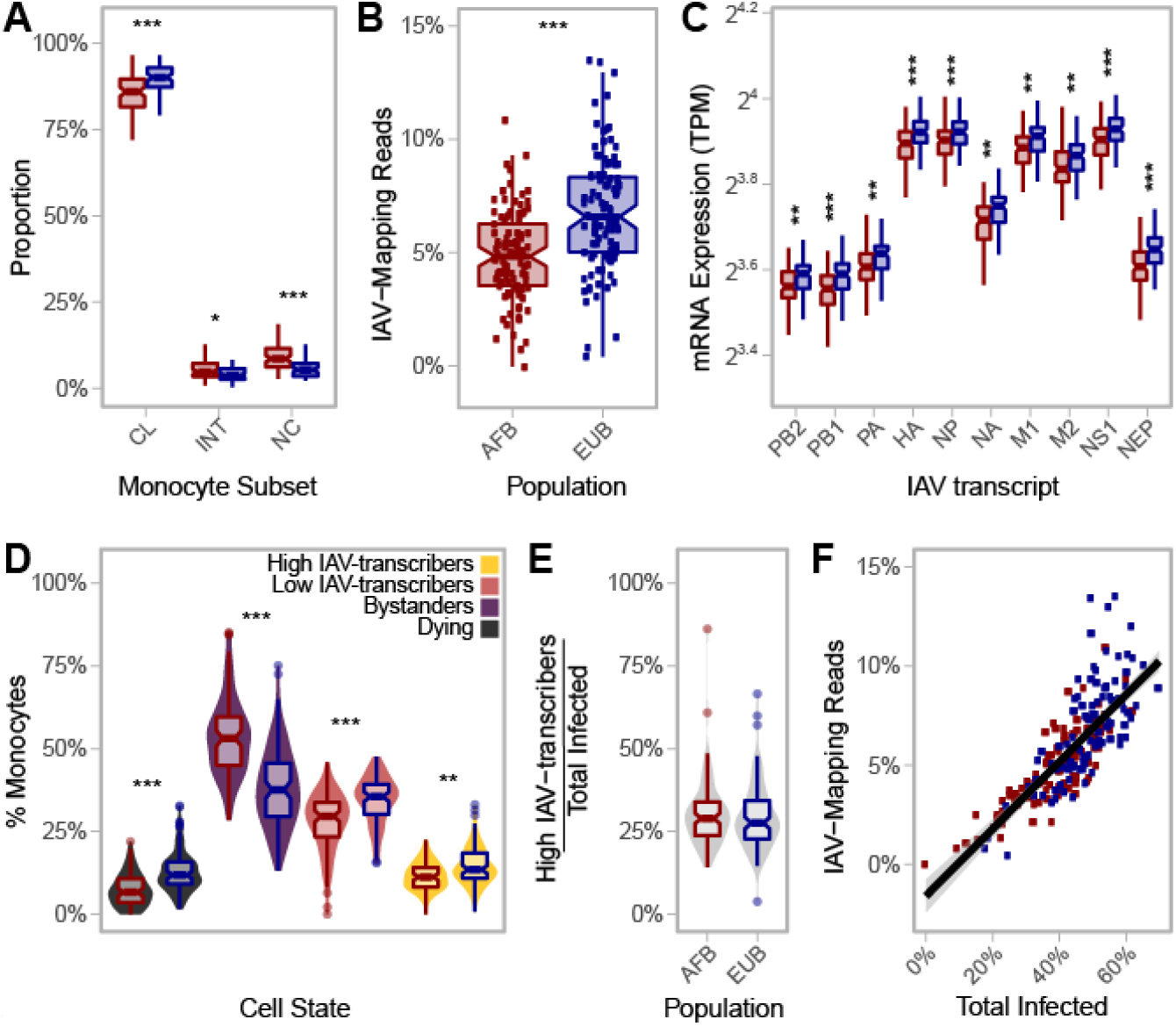
African-Ancestry Individuals Display Increased Number of *CD16*^*+*^ Cells and Lower Susceptibility to IAV Infection. (***A***) Variation in the number of classical (CD14^++^CD16^-^), intermediate (CD14^++^CD16^+^), and nonclassical (CD14^+^CD16^++^) monocytes across African- and European-ancestry individuals following CD14^+^ selection from PBMCs (*n*_AFB_=89, *n*_EUB_=85). Colors reflect population (AFB in red and EUB in blue). All three subsets are significantly different between populations (*p*-value<0.01). (***B***) Inter-and intra-population variation in the percentage of RNA-seq reads mapping to the IAV genome (*n*_AFB_=100, *n*_EUB_=99). Colors reflect population (AFB in red and EUB in blue). The percentage of RNA-seq reads mapping to the IAV genome is significantly higher in European-ancestry individuals relative to African-ancestry individuals (*p*-value=5.3×10^−8^). (***C***) Inter- and intra-population variation in viral mRNA expression at 6hpi (*n*_AFB_=100, *n*_EUB_=99). Expression levels for each of the 10 primary transcripts of IAV are plotted. Colors reflect population (AFB in red and EUB in blue). All IAV transcripts are significantly higher in European-ancestry individuals on average (*p*-value<0.001). (***D***) Estimated distribution of the percentage of cells from each cell state in the bulk RNA-seq data (*n*_AFB_ =100, *n*_EUB_=99). Fill colors reflect cell state assignments, while outlines of boxplots reflect population (AFB in red and EUB in blue). (***E***) Distribution of the percentage of high IAV-transcribers among infected cells, stratified by population. One individual with no infected cell was excluded (*n*_AFB_=99, *n*_EUB_=99). (***F***) Percentage of RNA-seq reads of viral origin as a function of the estimated proportion of infected cells (*n*_AFB_=100, *n*_EUB_=99), colored by population (AFB in red and EUB in blue). ***A* and *C*:** Outlier points are not displayed. Abbreviations: African-ancestry individuals from Belgium (AFB), European-ancestry individuals from Belgium (EUB), transcripts per million (TPM), influenza A virus (IAV), mean fluorescent intensity (MFI), classical (CL), intermediate (INT), and nonclassical (NC). * *p*-value < 0.01; ** *p*-value < 0.001; *** *p*-value < 0.0001

To test this hypothesis, we mapped the bulk RNA-seq profiles collected 6hpi challenge with IAV for the 200 individuals to a combined human-IAV reference. Excluding 1 sample with low quality RNAs, we found that 0.02-13.5% of RNA-seq reads from each sample were of viral origin (**Fig. 5*B***). Reassuringly, these percentages correlated with IAV mRNA levels estimated from the single cell experiment across all time points for the eight donors used in the present study (Pearson r > 0.84, *p*-value < 8.9×10^−3^), with the strongest correlation being observed at the peak of viral transcription (T_4_) (Pearson r = 0.97, *p*-value=5.1×10^−5^). These observations indicate that *ex vivo* cellular susceptibility is highly reproducible among individuals, even across different experimental protocols and technologies. Among the 199 bulk profiles, AFB and EUB samples presented overlapping but significantly shifted distributions of total IAV-mapping reads (**Fig. 5*B***, 4.9% vs. 6.8% of reads, respectively, Wilcoxon *p*-value = 5.3×10^−8^), and of each of the 10 primary viral transcripts (**Fig. 5*C***, Wilcoxon *p*-value < 5.5×10^−4^).

Using the transcriptional profiles obtained from the scRNA-seq data at T_6_, we estimated the proportion of reads coming from each inferred cell state in these bulk RNA-seq profiles (**Fig. 5*D* and *5E*; *SI Appendix*, Note S2 and Fig. S8*A***). We found that, on average, AFB monocytes were more resistant to IAV infection than EUB monocytes (39.2% vs. 48.9% infected, respectively, Wilcoxon *p*-value = 5.3×10^−10^). Differences in the estimated percentage of infected cells alone explained 63% of the inter-individual variability in viral mRNA levels (**Fig. 5*F***), and was sufficient to account for the observed difference in viral mRNA levels between AFB and EUB individuals (*p*-value=0.16 after adjusting on infected cells, compared to *p*-value=5.3×10^−8^ without adjustment). Nonetheless, variation in the percentage of high/low transcribers among infected cells accounted for an additional 19% of variance in viral mRNA expression (***SI Appendix*, Note S2 and Fig. S8*B***). Finally, the ratio of *CD16*^*+*^/*CD16*^*-*^ cells negatively correlated with the percentage of infected cells, albeit weakly (−0.27, *p*-value = 0.0165 adjusted on population). Altogether, these results show that population differences in viral mRNA levels are primarily driven by the overall proportion of cells that will ultimately become infected, with only a fraction of the differences being attributable to the different proportions of *CD16*^*+/-*^ subsets observed in individuals of African and European ancestry.

## Discussion

We performed scRNA-seq on primary monocytes, before and after *ex vivo* IAV-challenge, to assess transcriptional differences between monocytes infected by IAV (i.e. infected) versus those activated only by exposure to secreted cytokines (i.e. bystanders), and to identify subset-specific responses of monocytes to viral challenge. We found that bystander cells display increased mRNA expression of ISGs relative to infected cells; yet, we additionally observed both an induction of ribosomal gene mRNA expression in IAV-transcribing cells and a down regulation of these genes in bystander cells at later time points. While the former is likely induced by the virus to enhance mRNA translation (33), the repression of ribosomal expression observed in bystander cells may reflect a host mechanism to contain infection by shutting down the translational machinery of neighboring cells. Interestingly, the interplay of ribosomal and ISG expression also distinguished infected cells into two distinct states (high and low IAV-transcribers), providing an explanation for the high cell-to-cell variation in IAV replication observed among circulating monocytes, which has also been documented in other cell types and during natural infection (34-41).

While these patterns are generally shared across *CD16*^*-*^ and *CD16*^*+*^ subsets, we found *CD16*^*+*^ cells to be slightly more resistant to infection. This is likely attributable to their higher absolute expression of some ISGs relative to *CD16*^*-*^ cells (independent of viral exposure), as well as their more robust upregulation of antiviral genes upon IAV-challenge, which we found to be driven by stronger activity of IRF transcription factors. Interestingly, *CD16*^*+*^ cells displayed transient mRNA expression of *IL6* and *TNF* upon viral exposure (both infected and bystander cells), two cytokines that have been widely implicated in cytokine storms (5). Collectively, these findings highlight the opposing roles of ISG and ribosomal gene mRNA expression on viral transcription, and reveal the stronger antiviral and pro-inflammatory potential of *CD16*^*+*^ monocyte subsets.

At the population level, we found that the ratio of *CD16*^*+*^/*CD16*^*-*^ at basal state was predictive of the percentage of monocytes that were susceptible to IAV infection, and observed that African-ancestry individuals harbored more *CD16*^*+*^ monocytes on average than European-ancestry individuals residing in the same city (Ghent, Belgium), consistent with previous observations (42). Independently of monocyte subset proportions, we identified that individuals presenting lower monocyte susceptibility to IAV had a higher basal activation of an IRF/STAT-driven antiviral program. These findings suggest that the fate of a monocyte hinges upon its basal activation state, and that the infection potential differs both within an individuals’ monocyte population, in part based on the differentiation status of the cell (i.e. *CD16*-positivity), but also between individuals, where a *CD16*^*-*^ cell from one individual may have a higher antiviral state than a *CD16*^*+*^ cell from another individual. This latter phenomenon likely reflects the influence of both genetic and non-heritable factors on transcriptional variation in immune genes (14, 15).

These inter- and intra-population differences are noteworthy in and of themselves, and raise questions about how such differences in susceptibility of monocytic cells to infection - and differences in proportions of monocyte subsets - may relate to viral disease.

African Americans are more often hospitalized than other self-defined ethnic groups by both influenza and COVID-19, even when adjusting for age and various social factors such as poverty and vaccination status (43, 44). Thus, should monocyte infection play a role in the severity of viral infections, our results would imply a paradox where higher infection and replication of IAV in circulating monocytes is associated with an advantage to fight viral infection *in vivo*. Such a paradox could possibly be explained by an enhanced antigen presenting capacity of infected cells, with higher infection of European-monocytes facilitating the activation of T cell antiviral responses (13, 45). In support of this hypothesis, we observe that infected and bystander cells display comparable induction of antigen-presentation genes relative to non-exposed cells. This suggests that infected cells maintain their ability to present viral antigens to the adaptive immune system, while actively producing these antigens. Independently, but not mutually exclusively, proportions of monocyte subsets in circulation may play a role in viral disease; of note, patients with severe influenza and COVID-19 harbor higher proportions of intermediate monocytes in peripheral blood than patients with mild disease (46, 47). Given our finding that *CD16*^*+*^ subsets are the main drivers of inflammatory cytokine gene expression such as *IL6* and *TNF*, and that African-ancestry individuals harbor a larger fraction of these monocyte subsets, it is tangible to conceive that monocyte subset composition prior to infection may influence disease outcome, and potentially serve as a biomarker.

## Materials and methods

### Experimental model and subjects

All individuals from this study were part of the EVOIMMUNOPOP cohort, which has been previously described (18). Briefly, 200 healthy male donors living in Belgium of self-reported African descent (AFB) or European descent (EUB) were recruited. Inclusion was restricted to nominally healthy individuals between 19 and 50 years of age at the time of sample collection. Serological testing was performed for all donors to exclude those with serological signs of past or ongoing infection with human immunodeficiency virus (HIV), hepatitis B virus (HBV) or hepatitis C virus (HCV).

### Single-cell analyses and RNA sequencing

For eight selected donors (4 individuals from each ancestry, selected from extremes of the first principal component of gene expression in our previous study of monocyte response to IAV-challenge (18)), 100×10^6^ PBMCs were thawed, washed twice and resuspended in complete medium: pre-warmed RPMI-1640 Glutamax medium, supplemented with 10% FCS and 1% penicillin/streptomycin (Cat# 15140-122, Life Technologies). Monocytes were then positively selected with magnetic CD14 microbeads, according to the manufacturer’s instructions (Cat#130-050-201, Miltenyi Biotec). The number of monocytes was determined with the Countless2 automated cell counter system (Cat# AMQAX1000, ThermoFisher Scientific) in the presence of trypan blue. For each donor, monocytes were seeded at 0.5×10^6^ monocytes per well on 24-well NUNC plates in 500 µL of complete media and allowed to rest for one hour at 37°C under 5% CO_2_. Five-hundred microliters of complete media (non-infected) or A/USSR/90/1977(H1N1) at a concentration of 1×10^6^ pfu/mL in complete media (IAV-challenged, MOI=1) were added to each sample.

Following one hour of staging at 4°C, plates were centrifuged at 1300 rpm for 10 minutes at 4°C, media was removed by pipette, and each well was washed with 1mL complete media. The spin was repeated, media removed by pipette, and samples were resuspended in 1mL pre-warmed complete media before being transferred to an incubator at 37°C under 5% CO_2_ to initiate infection (T_0_).

At each time point (T_0_, T_2_, T_4_, T_6_, and T_8_), samples were mixed by pipetting and transferred to Eppendorf tubes. Wells were washed with 300uL of PBS + 0.04% BSA and transferred to the same tubes. Collection tubes were centrifuged at 1300 rpm for 10 minutes, media was removed and replaced with 1mL PBS + 0.04% BSA and an aliquot of 10µL was taken to count each sample on a Countless2 automated cell counter system, before repeating the centrifugations. Individual samples were adjusted to 2×10^6^ live cells/mL.

Samples were multiplexed for running on the 10X Chromium (Cat# 120223 & 1000074, 10X Genomics) by mixing equal proportions from 6-8 samples in a manner that balanced conditions and allowed us to assess for batch effects across lanes (***SI Appendix*, Table S1**). Multiplexed samples were counted with the Countless2 automated cell counter system and adjusted to target recovery of 10,000 cells per reaction of the Chromium Single Cell 3’ Reagent Kits v3 (Cat# 1000092 & 1000078, 10X Genomics) assuming a recovery rate of 50%. GEM Generation & Barcoding, Post GEM-RT Cleanup & cDNA Amplification, and 3’ Gene Expression Library Construction were performed as per manufacturer’s instructions (48). All 13 libraries were mixed prior to sequencing across 13 different lanes from an Illumina HiSeq X (28bp barcode + 91bp insert – target 400 M reads pairs per lane), leading to a total of 5.3 billon reads.

#### Sample Genotyping

Genotyping data [accession EGAS00001001895] were obtained for all 200 individuals from the EvoImmunoPop cohort based on both Illumina HumanOmni5-Quad BeadChips and whole-exome sequencing with the Nextera Rapid Capture Expanded Exome kit (18). The 3,782,260 SNPs obtained after stringent quality control were then used for imputation, based on the 1,000 Genomes Project imputation reference panel (Phase 1 v3.2010/11/23) (49), leading to a final set of 19,619,457 high-quality SNPs, of which 7,766,248 SNPs had a MAF ≥5% in our cohort.

#### Processing of scRNA-seq data

Basic pre-processing of the sequencing data was performed with CellRanger v3.0.2 (50), including the *mkfastq, count*, and *aggr* commands. Default parameters and our combined human-IAV reference were used, and batch correction was disabled in the *aggr* command. Cell-containing droplets (*n*=132,130) were traced back to individual donors using two independent methods, Demuxlet and SoupOrCell, which capitalize on genetic variation in the sequencing reads (20, 51). Barcodes with ambiguous and/or non-concordant calls between the two programs were used to establish suitable QC metrics. We found that barcodes deemed as *doublets* (i.e. the droplet contained two or more cells originating from different donors) were more likely to be nearest-neighbors in a *knn*-graph with other *doublets* than assigned *singlets*. We used this feature to identify droplets presumed to contain two or more cells originating from the same donor; barcodes with > 5 *doublets* as nearest-neighbors were excluded from further analysis (***SI Appendix*, Fig. S1 *A* and *B***). Additionally, droplets containing low-quality cells (i.e. damaged, dying) were excluded using the following thresholds: < 1500 total counts, < 500 genes, or > 50% mitochondrial gene content (***SI Appendix*, Fig. S1*C***). This QC resulted in a final data set of 96,386 single cells.

Transcriptomes (i.e. counts) were adjusted for the presence of ambient RNA with SoupX, (https://github.com/constantAmateur/SoupX/, accessed November 28, 2019) (21), using estimated contamination fractions (per 10X library) from SoupOrCell (51). SoupX-adjusted counts were normalized using pool-based size factors followed by deconvolution as implemented in the scran R package (52). Feature selection was performed by (i) constructing a mean-variance trend in the log-counts and retaining genes found to exhibit more variation than expected assuming Poisson-distributed technical noise, as implemented in the makeTechTrend and TrendVar functions from package scran (52), and (ii) selecting genes expressed in at least 25 cells (*n*=22,603). The first 10 PCs of the data were retained for data visualization and clustering analyses. Graph-based clustering was performed by building the shared nearest-neighbor graph with the buildSNNGraph function from scran (52) using a series of *k* values, and cell clusters were defined with the igraph Walktrap algorithm (53). Similar clustering results were obtained based on the *knn*-graphs generated using *k*=25, 50, 75, and 100, and *k*=25 was used for all downstream analyses (***SI Appendix*, Fig. S2*A***). Cell types were predicted using SingleR and the built-in BlueprintEncodeData reference (54). Based on the clustering and cell-type predictions, we removed cells belonging to clusters associated with lymphoid cell types or low QC metrics from downstream analyses (***SI Appendix*, Fig. S2 *B* and *C***).

### Accounting for ambient RNA contamination in scRNA-Seq Data and assigning cell states

Droplet-based scRNA-seq methods capture ambient mRNAs present in the cell suspension in addition to cell specific mRNAs. To estimate which cells in our experiment were genuinely expressing mRNAs for *CD14, FCGR3A (CD16)*, and those originating from the virus, we implemented a two-step strategy utilizing the estimateNonExpressionCells function of the SoupX package (21). This function estimates whether each cell contains significantly more counts of a provided gene-set than would be expected under a Poisson model, given the estimated ambient RNA from its library of origin and the maximum contamination fraction. First, we used the viral genes to estimate the true maximum contamination fraction, based on the assumption that cells from the non-infected state should only contain viral reads from ambient mRNA captured in their droplets. To do so, we modified the estimateNonExpressionCells function to return *p*-values, and performed the test on each of our 13 libraries with a range of maximum contamination values from 1-50% (step of 1%) using the viral genes. We then computed FDR adjusted *p*-values for each maximum contamination value on the 88,559 high-quality, single monocytes. The number of non-simulated cells deemed to significantly express IAV transcripts (FDR<0.01) was used as a proxy for false positives. In examining the relationship between this number and the number of IAV-challenged cells found to significantly express viral transcripts at FDR<0.01 (**Fig. *1C***), we found that a maximum contamination fraction of 10% resulted in a 1% false positive rate (defined as the percentage of non-infected cells from T_2_-T_8_ that were deemed to significantly express IAV transcripts). This parameter value was then used to correct for contamination from ambient for all genes considered (*CD14, FCGR3A* and IAV transcripts).

### Assigning cell states and investigating sources of variability in IAV levels

We used a maximum contamination fraction of 10% to test for significant expression of IAV transcripts in each cell (**Fig. 1 *C-E***). IAV-challenged cells that contained a significant amount of IAV transcripts were considered as infected, while the others were deemed bystanders. To distinguish low from high IAV-transcribing cells, a Gaussian mixture model was fitted to the total percentage of viral mRNAs per cell across all infected cells, using the normalmixEM function from mixtools R package with *k*=2 (55). Each cell was assigned to the cluster with the highest posterior probability, and the cluster of cells with higher IAV content was annotated as high IAV-transcribing.

### Characterizing monocyte subsets and transcriptional profiles from scRNA-seq data

Principal components analysis of 6,601 cells at T_0_ was used to order monocytes along a differentiation axis separating *CD14*^*+*^ cells from *CD16*^*+*^ cells. We then computed the average percentage of classical and nonclassical monocytes obtained by FACS across the eight donors, weighting each individual by the number of high-quality cells in the scRNA-seq data at T_0_. Based on these percentages (87.1% for classical and 7.6% for nonclassical), we annotated the monocytes on each side of the differentiation axis as classical and nonclassical, respectively, with the remaining 5.3% of monocytes being annotated as intermediates. Validity of our approach was confirmed by correlating the proportion of monocytes assigned to each subset across the eight donors, with the percentage of classical, intermediate and nonclassical monocytes estimated by FACS.

Differential expression between subsets was assessed for the 4,859 genes expressed at an average logCount > 0.1 in any of the 3 subsets. Specifically, Wilcoxon rank tests were implemented in the scran package (52), using the findMarkers function and blocking on donor. We considered genes to be differentially expressed (DE) between monocyte subsets when gene expression was significant at an FDR≤1% and log_2_FC>0.2. The 848 genes that differed between classical (CL) and nonclassical (NC) monocyte subsets were classified according to their behavior in intermediate monocytes (INT). They were either deemed ‘similar to classical’ (DE between INT and NC, but not between INT and CL), ‘similar to nonclassical’ (DE between INT and CL, but not between INT and NC), or ‘intermediate’ (all other cases).

At later time points, comparisons between *CD16*^*+*^ and *CD16*^*-*^ monocytes subsets were done based on 5,681 genes expressed with a normalized log count > 0.1 on average in either subset, in at least one condition and time point. For each subset, log_2_ fold change in gene expression relative to T_0_ were correlated across times points. Differential expression between *CD16*^*+*^ and *CD16*^*-*^ cells was assessed with findMarkers (52), based on Wilcoxon rank tests and blocking on donors, time points and condition. Again, an FDR≤1% and log_2_FC>0.2 were required to define differentially expressed genes. To assess how CD16^+/-^ status alters the infection of monocytes by IAV, we used logistic regression to model bystander/infected status as a function of *CD16*^*+/-*^ status, while adjusting on donor, and time point (as factors).

### Characterizing subset-specific responses to IAV challenge

To allow comparison between responses of *CD16*^*+*^ and *CD16*^*-*^ monocytes, 100 cells were subsampled from each subset and cell state, and at each time point. When subsampling, we ensured balanced representation of all donors across each monocyte subset and cell state, by using sampling weights that were inversely proportional to each donor representation in the original dataset. After sampling, a total of 6,669 genes with normalized log counts >0.1 on average in at least one group (cell-state x subset x time point) was selected for further analyses. For each monocyte subset, differences in expression between cell states (unexposed, bystander, infected) as well as between high- and low-IAV transcribing infected cells were performed using the findMarkers function from the scran package (52) and blocked on time-point. For each comparison, genes were considered to be differentially expressed between cell states when gene expression was significant at an FDR=1% (Wilcoxon rank tests) and the log_2_ fold change was > 0.2. In addition, for each comparison between cell states, we tested for differences in response between subsets using a linear model of the form:

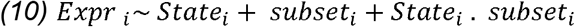

Where *Expri_i_* is the expression of the gene being tested in cell *i, state_i_* is an indicator variable that distinguishes the two cell states being compared (e.g. unexposed and bystander), and *subset_i_* is an indicator variable that reflects the *CD16*^*+/-*^ status of cell *i*. The 335 genes with significant interactions at a 1% FDR (for unexposed-bystander and unexposed-infected comparisons) were clustered using the hclust R function with method ‘Ward.D2’. DynamicTreeCut algorithm (56) was used to identify eight major patterns of response to IAV.

### Transcription factor enrichment analyses

To estimate Transcription Factor (TF) activity and define TF-targets relationships, we ran the R SCENIC pipeline (29) on the expression matrix (pre-normalization) on a random subsample of 4800 cells (100 cells from each donor at each time point and each condition, pre-exclusion of dying and contaminant cells) with default parameters. For each gene, motif-enrichment was considered for either *cis*-regulatory regions located <10kb from the TSS (distal regulatory elements), or between 500 bp upstream and 100 bp downstream of the promoter (proximal regulatory elements). To do so, motif-enrichment scores for all human genes (hg38 build, refseq_r80), were retrieved from https://resources.aertslab.org/cistarget and used as input for the Rcistarget package (29).

Sets of high-confidence targets for the 113 TFs whose activity could be quantified by SCENIC were then extracted and used for enrichment analysis. For each gene module, TF enrichment was assessed using a Fisher’s exact test with the 6,669 expressed genes as background (**Dataset 3**). Resulting *p*-values were adjusted using a global Benjamini-Hochberg correction for all eight modules and 113 TFs.

For each TF, with its targets enriched among one of the eight modules, TF activity inferred by SCENIC was used to test for subset-specific activity using a linear model of the form:

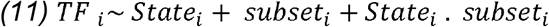

where *TF_i_* is the activity of the TF being tested in cell *i, statei_i_* is an indicator variable that distinguishes the two cell states being compared (e.g. unexposed and bystander), and *subset_i_* is an indicator variable that reflects the *CD16*^*+/-*^ status of cell. Average TF activity was then computed for each cell state, subset and time point, and correlated with gene expression of the associated module, to assess the link between TF activation and the TF-target enriched modules.

### Association of the outcome of IAV infection using basal gene expression

For each of the three monocyte subsets detected at basal state, Kruskall-wallis test was used to search for genes whose expression levels significantly differ across donors. Within each monocyte subset, we then computed the average expression of each gene for all eight donors and correlated it with the percentage of infected cells at 4hpi. Genes that differed in expression between donors (FDR≤1%), and passed a nominal *p*-value threshold of 0.01 for association with IAV levels in any of the 3 subsets, were selected for downstream enrichment analyses. For genes nominally correlated with viral mRNA levels, TF enrichment was assessed as previously using a Fisher’s exact test with all 4,859 genes expressed at T_0_ as background (**Dataset S4**), and Benjamini-Hochberg correction for all 113 TFs was applied.

### Gene Ontology enrichment analyses

All Gene Ontology (GO) enrichment analyses were performed with GOSeq package using default settings (57). Background gene sets consisted of all genes that had average log-normalized expression value > 0.1 in at least one of the groupings being examined, and are described in the text. Only enrichments significant at FDR≤5% are reported.

### Pseudo-bulk estimates from scRNA-seq data

Pseudo bulk estimates of IAV mRNA levels were computed by measuring, for each donor and time point, the mean percentage of reads of viral origin across all cells from the sample. At each time point, we then used a Pearson’s correlation test to compare pseudo-bulk estimates for the 8 donors with IAV mRNA levels obtained in bulk data at 6hpi.

### Monocyte subset characterization of EVOIMMUNOPOP samples via FACS

For 174 of the 200 EVOIMMUNOPOP donors, proportions of classical, intermediate and nonclassical monocytes were determined based on a fraction of 10^5^ CD14^+^ positively-selected monocytes, stained according to the manufacturer’s instructions, with fluorescent APC-conjugated anti-CD14 and PE-conjugated anti-CD16 antibodies (Cat#130-091-243 and Cat #130-091-245, respectively, Miltenyi Biotec). Samples were then analyzed on a MACSQuant Analyzer 10 benchtop flow cytometer (Miltenyi Biotec).

### Quantification of canonical monocyte subsets in EVOIMMUNOPOP samples

FlowJo v10.6.1 software (58) was used with the gating strategy depicted in ***SI Appendix*, Fig. S1** to quantify monocyte subsets for 174 EVOIMMUNOPOP donors. Population-level differences in proportion of canonical monocyte subsets were assessed using Wilcoxon Rank tests.

Correlation of the ratio of CD16^+^ to CD16^-^ cells with IAV mRNA levels was assessed using a linear model of the form

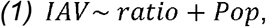

where ‘*IAV’* are IAV mRNA levels, ‘*ratio*’ is the percentage of CD16^+^ monocytes (non classical+intermediates) divided by the percentage of CD16^-^ monocytes (classical), and ‘*Pop’* is and indicator variable separating AFB from EUB individuals.

### Analysis of bulk RNA-seq profiles from the EVOIMMUNOPOP cohort

A combined human-IAV reference was generated by concatenation of the primary human genome assembly (GRCh38) with the 8 segments of the human influenza A virus (IAV) A/USSR/90/1977(H1N1) genome (accession numbers CY010372-CY010379). Comprehensive human gene annotation was obtained from GENCODE (release 27) and merged with the 12 known transcripts of A/USSR/90/1977(H1N1). RNA-seq reads (FASTQs) for all 970 samples that passed quality control in our previous study (18) [accession EGAS00001001895] were mapped to the combined reference with the STAR aligner (v.2.5.0a) (59) and assessed for quality with QualiMap ‘bamqc’ and ‘rnaseq’ (60, 61). Expression of viral mRNAs was measured as the percentage of uniquely-mapped reads aligning to the IAV genome. Reassuringly, the mean percentage of RNA-seq reads among samples from the IAV-challenged condition was 5.86% versus <0.01% in the other four conditions. Comparison of the percentage of IAV reads between populations was done using a Wilcoxon rank test. StringTie (v.1.3.3) (62) was used to quantify expression levels in transcripts per million mapped reads (TPM) for each annotated transcript. Gene expression data were filtered to remove genes with little evidence of activation (mean zTPM score < -3) (63) in any of the 5 conditions, and their quality was checked by principal component analysis (PCA). As GC content, 5′/3′ bias, date of the experiment and library batch were previously determined to be the strongest confounding factors on transcript expression (18), we corrected the data for these factors. First, we adjusted the data for GC content and 5′/3′ bias using linear models. Then, we imputed missing values by k-nearest neighbor imputation and adjusted for experiment date and library batch by sequentially running ComBat (64) for each batch effect, with condition and population as covariates. After batch effect correction, only IAV-stimulated samples were kept for downstream analyses.

### Cell states deconvolution from bulk RNA sequencing

To assess the percentage of total transcripts that originate from each cell state across the 199 IAV-challenged samples, we pooled cells from T_6_ into 3 groups, based on their assigned cell-state (bystander, infected: high and low IAV-transcribing) and to which we added a 4^th^ group containing all singlets that were either (i) assigned to cluster numbers 3, 8, 10, and 11 (dying cells) or (ii) discarded based on their high mitochondrial content or low read counts (dead cells). We then estimated pseudo-bulk profiles for each group by summing UMIs across all cells and computing the number of UMIs associated to each gene per million of sequenced UMIs. TPM profiles obtained from bulk data were then normalized to improve comparison with pseudo-bulk. Specifically, we first computed a global pseudo-bulk profile of the entire single cell dataset as the average of the pseudo bulk profiles from the 4 cell states (bystander, infected: high and low IAV-transcribing, or dying/dead), weighted by the percentage of UMIs they contribute to the overall pool of cells. To account for the difference in how gene expression is quantified between the two methods (3’ end counts for scRNA-seq and full-length gene coverage for bulk RNA-seq), we computed for each gene i a normalization factor *s_i_* given by

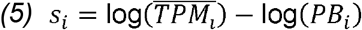

where *PB_i_* is the number of UMI per million for gene *i* in the global pseudo-bulk profile, and 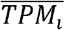 is the average expression of the gene *i* in the 199 IAV-stimulated samples from the bulk RNA-seq data. For each gene, *s_i_* was then subtracted from the log transformed TPM to yield a normalized TPM profiles. We next applied DeconRNAseq (65) to the normalized log TPM profiles from all individuals, using the log-transformed pseudo bulk profiles from the 4 cell states as a basis for deconvolution. Quality of the deconvolution was assessed using leave-one-out cross validation, based on the eight individuals for whom we had scRNA-seq data. Specifically, for each of these eight individuals, bulk mRNAs were decomposed using pseudo-bulk profiles recomputed based on the seven other individuals. The resulting proportions were then compared with the percentage of UMIs that originate in each cell-state in the scRNA-seq to assess the quality of the deconvolution. Comparisons between populations were performed using Wilcoxon rank tests.

The effect of the percentage of infected cells and percentage of high IAV-transcribing cells among infected cells on the total IAV mRNA levels were assessed by modeling

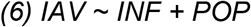

and

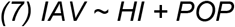

where *IAV* are the IAV mRNA levels across the 199 bulk mRNA samples, *INF* and *HI* are respectively the percentage of infected cells and the percentage of high IAV-transcribing cells among infected cells that we estimated from the deconvolution, and *POP* is a factor variable reflection the population (EUB or AFB). The fraction η of population differences attributable to difference in rate of infection was estimating by comparing model (6) with model (8) below

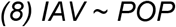

and computing 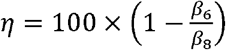, with β_*i*_ the effect of population on IAV levels in model (i). To assess how the contribution of the percentage of high IAV transcribing cells to total IAV mRNA levels differed between populations, we used a linear model of the form

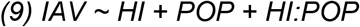

and tested for significant effect of the interaction term *HI:POP* on *IAV* mRNA levels.

## Data availability

The bulk RNA-seq data used in this study are available at the European Genome Phenome archive under accession number [EGAS00001001895]. The single cell RNA-seq data generated during this study are available at the European Genome Phenome archive under accession number [EGAS00001005000]. Code generated as part of this study is available on Github (https://github.com/h-e-g/PopDiff_MonocyteIAV).

## Acknowledgments

This work was supported by the *Institut Pasteur*, the *Collège de France*, the French Government’s Investissement d’Avenir program, *Laboratoires d’Excellence* “Integrative Biology of Emerging Infectious Diseases” (ANR-10-LABX-62-IBEID) and “Milieu Intérieur” (ANR-10-LABX-69-01), the *Fondation de France* (n°00106080), and the *Fondation pour la Recherche Médicale* (Equipe FRM DEQ20180339214). M.B.O was supported by a *European Molecular Biology Organization* long-term fellowship (ALTF 229-2017). We also wish to thank Tzachi Hagai and John Marioni for input and feedback, and Sylvie van Der Werf and Vincent Enouf for providing resources and protocols relating to influenza viruses.

## References

1. V. A. Ryabkova, L. P. Churilov, Y. Shoenfeld, Influenza infection, SARS, MERS and COVID-19: Cytokine storm - The common denominator and the lessons to be learned. Clinical immunology (Orlando, Fla.) 223, 108652 (2021).

2. F. Krammer et al., Influenza. Nat Rev Dis Primers 4, 3 (2018).

3. Q. Zhang et al., Life-Threatening COVID-19: Defective Interferons Unleash Excessive Inflammation. Med (N Y) 1, 14–20 (2020).

4. A. A. Stegelmeier et al., Myeloid Cells during Viral Infections and Inflammation. Viruses 11 (2019).

5. D. C. Fajgenbaum, C. H. June, Cytokine Storm. N Engl J Med 383, 2255–2273 (2020).

6. C. Guo et al., Single-cell analysis of two severe COVID-19 patients reveals a monocyte-associated and tocilizumab-responding cytokine storm. Nature communications 11, 3924 (2020).

7. R. Alon et al., Leukocyte trafficking to the lungs and beyond: lessons from influenza for COVID-19. Nat Rev Immunol 21, 49–64 (2021).

8. L. Ziegler-Heitbrock et al., Nomenclature of monocytes and dendritic cells in blood. Blood 116, e74–80 (2010).

9. F. Geissmann, S. Jung, D. R. Littman, Blood monocytes consist of two principal subsets with distinct migratory properties. Immunity 19, 71–82 (2003).

10. C. Auffray et al., Monitoring of blood vessels and tissues by a population of monocytes with patrolling behavior. Science 317, 666–670 (2007).

11. L. Ziegler-Heitbrock, The CD14+ CD16+ blood monocytes: their role in infection and inflammation. J Leukoc Biol 81, 584–592 (2007).

12. M. A. Hoeve, A. A. Nash, D. Jackson, R. E. Randall, I. Dransfield, Influenza virus A infection of human monocyte and macrophage subpopulations reveals increased susceptibility associated with cell differentiation. PLOS ONE 7, e29443 (2012).

13. W. Hou et al., Viral infection triggers rapid differentiation of human blood monocytes into dendritic cells. Blood 119, 3128–3131 (2012).

14. B. Piasecka et al., Distinctive roles of age, sex, and genetics in shaping transcriptional variation of human immune responses to microbial challenges. Proceedings of the National Academy of Sciences of the United States of America 115, E488–E497 (2018).

15. P. Brodin et al., Variation in the human immune system is largely driven by non-heritable influences. Cell 160, 37–47 (2015).

16. W. J. Astle et al., The Allelic Landscape of Human Blood Cell Trait Variation and Links to Common Complex Disease. Cell 167, 1415–1429 e1419 (2016).

17. E. Patin et al., Natural variation in the parameters of innate immune cells is preferentially driven by genetic factors. Nat Immunol 19, 302–314 (2018).

18. H. Quach et al., Genetic Adaptation and Neandertal Admixture Shaped the Immune System of Human Populations. Cell 167, 643–656 e617 (2016).

19. K. G. Ouwens et al., A characterization of cis-and trans-heritability of RNA-Seq-based gene expression. Eur J Hum Genet 28, 253–263 (2020).

20. H. M. Kang et al., Multiplexed droplet single-cell RNA-sequencing using natural genetic variation. Nature biotechnology 36, 89–94 (2018).

21. M. D. Young, S. Behjati, SoupX removes ambient RNA contamination from droplet-based single-cell RNA sequencing data. Gigascience 9 (2020).

22. K. L. Wong et al., Gene expression profiling reveals the defining features of the classical, intermediate, and nonclassical human monocyte subsets. Blood 118, e16–31 (2011).

23. V. Segura et al., In-Depth Proteomic Characterization of Classical and Non-Classical Monocyte Subsets. Proteomes 6 (2018).

24. C. Schmidl et al., Transcription and enhancer profiling in human monocyte subsets. Blood 123, e90–99 (2014).

25. A. Bercovich-Kinori et al., A systematic view on influenza induced host shutoff. eLife 5 (2016).

26. H. M. Machkovech, J. D. Bloom, A. R. Subramaniam, Comprehensive profiling of translation initiation in influenza virus infected cells. PLoS pathogens 15, e1007518 (2019).

27. S. Li, Regulation of Ribosomal Proteins on Viral Infection. Cells 8 (2019).

28. J. Cros et al., Human CD14dim monocytes patrol and sense nucleic acids and viruses via TLR7 and TLR8 receptors. Immunity 33, 375–386 (2010).

29. S. Aibar et al., SCENIC: single-cell regulatory network inference and clustering. Nature methods 14, 1083–1086 (2017).

30. E. K. Allen et al., SNP-mediated disruption of CTCF binding at the IFITM3 promoter is associated with risk of severe influenza in humans. Nature Medicine 23, 975 (2017).

31. Q. Zhang, Human genetics of life-threatening influenza pneumonitis. Human genetics 139, 941–948 (2020).

32. M. J. Ciancanelli et al., Infectious disease. Life-threatening influenza and impaired interferon amplification in human IRF7 deficiency. Science 348, 448–453 (2015).

33. B. Panthu et al., The NS1 Protein from Influenza Virus Stimulates Translation Initiation by Enhancing Ribosome Recruitment to mRNAs. J Mol Biol 429, 3334–3352 (2017).

34. C. Wang et al., Cell-to-Cell Variation in Defective Virus Expression and Effects on Host Responses during Influenza Virus Infection. mBio 11 (2020).

35. Y. Steuerman et al., Dissection of Influenza Infection In Vivo by Single-Cell RNA Sequencing. Cell systems 6, 679–691.e674 (2018).

36. A. B. Russell, C. Trapnell, J. D. Bloom, Extreme heterogeneity of influenza virus infection in single cells. eLife 7 (2018).

37. J. Sun et al., Single cell heterogeneity in influenza A virus gene expression shapes the innate antiviral response to infection. PLoS pathogens 16, e1008671 (2020).

38. I. Ramos et al., Innate Immune Response to Influenza Virus at Single-Cell Resolution in Human Epithelial Cells Revealed Paracrine Induction of Interferon Lambda 1. Journal of virology 93 (2019).

39. A. B. Russell, E. Elshina, J. R. Kowalsky, A. J. W. Te Velthuis, J. D. Bloom, Single-Cell Virus Sequencing of Influenza Infections That Trigger Innate Immunity. Journal of virology 93 (2019).

40. E. Kudo et al., Low ambient humidity impairs barrier function and innate resistance against influenza infection. Proceedings of the National Academy of Sciences of the United States of America 116, 10905–10910 (2019).

41. Y. Cao et al., Single-cell analysis of upper airway cells reveals host-viral dynamics in influenza infected adults. bioRxiv 10.1101/2020.04.15.042978, 2020.2004.2015.042978 (2020).

42. L. J. Appleby et al., Sources of heterogeneity in human monocyte subsets. Immunol Lett 152, 32–41 (2013).

43. R. Chandrasekhar et al., Social determinants of influenza hospitalization in the United States. Influenza Other Respir Viruses 11, 479–488 (2017).

44. J. L. Hadler et al., Influenza-Related Hospitalizations and Poverty Levels - United States, 2010-2012. MMWR Morb Mortal Wkly Rep 65, 101–105 (2016).

45. N. J. Roberts, Jr., Diverse and Unexpected Roles of Human Monocytes/Macrophages in the Immune Response to Influenza Virus. Viruses 12 (2020).

46. S. L. Cole et al., M1-like monocytes are a major immunological determinant of severity in previously healthy adults with life-threatening influenza. JCI Insight 2, e91868 (2017).

47. Y. H. Zhou et al., Pathogenic T cells and inflammatory monocytes incite inflammatory storm in severe COVID-19 patients. Natl Sci Rev 10.1093/nsr/nwaa041, nwaa041 (2020).

48. 10xGenomics, Chromium Single Cell 3’ Reagent Kits User Guide (v3 Chemistry). (2020).

49. C. Genomes Project et al., An integrated map of genetic variation from 1,092 human genomes. Nature 491, 56–65 (2012).

50. G. X. Zheng et al., Massively parallel digital transcriptional profiling of single cells. Nature communications 8, 14049 (2017).

51. H. Heaton et al., Souporcell: robust clustering of single-cell RNA-seq data by genotype without reference genotypes. Nature methods 17, 615–620 (2020).

52. A. T. Lun, D. J. McCarthy, J. C. Marioni, A step-by-step workflow for low-level analysis of single-cell RNA-seq data with Bioconductor. F1000Res 5, 2122 (2016).

53. P. Pons, M. Latapy (2005) Computing Communities in Large Networks Using Random Walks. in Computer and Information Sciences - ISCIS 2005, eds p. Yolum, T. Güngör, F. Gürgen, C. Özturan (Springer Berlin Heidelberg, Berlin, Heidelberg), pp 284–293.

54. D. Aran et al., Reference-based analysis of lung single-cell sequencing reveals a transitional profibrotic macrophage. Nat Immunol 20, 163–172 (2019).

55. T. Benaglia, D. Chauveau, D. R. Hunter, D. S. Young, mixtools: An R Package for Analyzing Mixture Models. Journal of Statistical Software; Vol 1, Issue 6 (2010) (2009).

56. P. Langfelder, B. Zhang, S. Horvath, Defining clusters from a hierarchical cluster tree: the Dynamic Tree Cut package for R. Bioinformatics 24, 719–720 (2008).

57. M. D. Young, M. J. Wakefield, G. K. Smyth, A. Oshlack, Gene ontology analysis for RNA-seq: accounting for selection bias. Genome Biol 11, R14 (2010).

58. D. a. C. Becton (2019) FlowJo™ Software (Ashland).

59. A. Dobin et al., STAR: ultrafast universal RNA-seq aligner. Bioinformatics 29, 15–21 (2013).

60. F. García-Alcalde et al., Qualimap: evaluating next-generation sequencing alignment data. Bioinformatics 28, 2678–2679 (2012).

61. K. Okonechnikov, A. Conesa, F. García-Alcalde, Qualimap 2: advanced multi-sample quality control for high-throughput sequencing data. Bioinformatics 32, 292–294 (2016).

62. M. Pertea et al., StringTie enables improved reconstruction of a transcriptome from RNA-seq reads. Nature biotechnology 33, 290–295 (2015).

63. T. Hart, H. K. Komori, S. LaMere, K. Podshivalova, D. R. Salomon, Finding the active genes in deep RNA-seq gene expression studies. BMC genomics 14, 778 (2013).

64. W. E. Johnson, C. Li, A. Rabinovic, Adjusting batch effects in microarray expression data using empirical Bayes methods. Biostatistics (Oxford, England) 8, 118–127 (2007).

65. T. Gong, J. D. Szustakowski, DeconRNASeq: a statistical framework for deconvolution of heterogeneous tissue samples based on mRNA-Seq data. Bioinformatics 29, 1083–1085 (2013).

