## Supplementary material for "Heterogeneity of monocyte subsets and susceptibility to influenza virus contribute to inter-population variability of protective immunity": SI Appendix

### Supplementary Notes

Note S1. scRNA-seq Accurately Identifies Classical, Intermediate, and Nonclassical Monocyte Subsets and their Functional Roles.

Principle component analysis (PCA) of transcriptional profiles of the 6,601 monocytes at T_0_ revealed a bimodal distribution of cells along PC1, which largely distinguished *CD14^+^* from *CD16^+^* cells (***SI Appendix*, Fig. S4*A***). We then binned cells into the three canonical monocyte subsets based on their PC1 values, with thresholds chosen to match the average proportions estimated by flow cytometry. To assess the performance of our approach, we measured how well we inferred the proportions of classical, intermediate, and nonclassical monocytes from the scRNA-seq data, and observed high concordance with those determined using flow cytometry (Pearson’s r = 0.88-0.97, *p*-values < 0.01).

To assess how the basal transcriptional profiles of canonical monocyte subsets differ, we focused on the 4,589 genes which were expressed, at a log normalized count > 0.1 on average, in at least one of the three monocyte subsets at T_0_ (**Dataset S1*A***). We found that 848 genes significantly differed between classical and nonclassical subsets (FDR<1%, log_2_FC>0.2), with 59% and 41% being upregulated in classical and nonclassical subsets, respectively (***SI Appendix*, Fig. S4*B***). Consistent with previous reports (1-3), classical monocytes were characterized by high expression of several proinflammatory *S100 Calcium Binding Proteins* (*S100A12*, *S100A9*, and *S100A8*) and the antimicrobial gene *Lysozyme* (*LYZ*)*,* contributing to sizable enrichments in the defense response to fungus (GO:0050832: OR=49.5, FDR=5.9x10^-4^) and antimicrobial humoral response (GO:0019730: OR=16.6, FDR=3.4x10^-5^) pathways. GO term enrichment analysis of genes highly expressed in the classical relative to the nonclassical subset (**Dataset S1*B***) also uncovered significant enrichment of genes implicated in tissue repair functions such as response to wounding (GO:0009611: OR=3.0, FDR=6.9x10^-7^), angiogenesis (GO:0001525: OR=2.2, FDR=1.9x10^-6^), and positive regulation of coagulation (GO:0050820; OR=16.5, FDR=4.2 x10^-3^). Conversely, the nonclassical subset was characterized by a strong over-expression of genes involved in cytoskeleton organization (GO:0032956: OR=4.1, FDR=4.3x10^-5^) and Fc-gamma receptor-mediated phagocytosis (GO:0038096: OR=5.7, FDR=9.6x10^-4^), as expected(1-3), but also of known regulators of lymphocytes proliferation (GO:0050670: OR=4.3, FDR=2.1x10^-4^) including the *Leukocyte specific transcript* *1* (*LST1*) and the *T-lymphocyte activation antigen CD86* (*CD86*)*.*

We then assessed the extent to which intermediate monocytes were related to both classical and nonclassical subsets. Among the 848 genes that were consistently differentially expressed between the donors’ classical and nonclassical monocytes (FDR<1%, log_2_FC>0.2), the intermediate subset displayed similar transcriptional levels to both classical (55 genes, e.g. *Carboxypeptidase Vitellogenic Like*, *CPVL*) and nonclassical (364 genes, e.g. *S100A12*) subsets for some genes, and intermediate levels for others (424 genes, e.g. *LYZ*) (***SI Appendix*, Fig. S4*C***). This supports to the notion that intermediate monocytes represent a transitionary state between classical and nonclassical subsets, and exhibit a transcriptional profile more closely related to that of the nonclassical population (1). Only 18 genes, five of which differed between classical and nonclassical subsets, were upregulated in the intermediate subset relative to both other subsets, at a log fold change ≥ 0.2 (designated as triangles in ***SI Appendix*, Fig. S4*B***). Among these 18 genes, we found 7 members of the major histocompatibility complex (MHC) class II protein complex, leading to enrichments of antigen processing and presentation pathways (e.g. GO:0019886; OR=112.1, FDR<1.1x10^-16^). Collectively, these results indicate that the monocyte subsets identified by scRNA-seq broadly recapitulate canonical monocyte populations classically defined via labeling for CD14 and CD16 cell surface antigens (FACS), and inform us on the functional impact of transcriptional heterogeneity between monocyte subsets.

1.	K. L. Wong *et al.*, Gene expression profiling reveals the defining features of the classical, intermediate, and nonclassical human monocyte subsets. *Blood* **118**, e16-31 (2011).

2.	V. Segura *et al.*, In-Depth Proteomic Characterization of Classical and Non-Classical Monocyte Subsets. *Proteomes* **6** (2018).

3.	C. Schmidl *et al.*, Transcription and enhancer profiling in human monocyte subsets. *Blood* **123**, e90-99 (2014).

Note S2. Deconvolution analysis reveals stronger impact of high IAV-transcribing infected cells on viral mRNA levels in European-ancestry individuals.

We first assessed the quality of our deconvolution method by cross-validation, comparing across our eight donors the estimated cell fractions obtained from deconvolution of the bulk RNA-seq profiles with the true proportions assessed through scRNA-seq (***SI Appendix*, Fig. S8*A***). We reliably captured inter-individual variation in the percentage of reads that originate from bystander (Pearson r = 0.97, p-value=4.2×10^-5^) and infected cells (Pearson r = 0.98, p-value=3.7×10^-5^), as well as, to a lesser extent, from dying/dead cells (Pearson r = 0.74, p-value=3.5×10^-2^). Applying the method to all 199 samples passing QC, we found that the percentage of reads from infected cells (both low and high IAV-transcribers) ranged from 0 to 70%, with this percentage being between 38 and 52% in more than half of the donors (**Fig. 5*D***). Our deconvolution method further allowed the separation of infected cells into low and high IAV-transcribers, although it was slightly less accurate at doing so (Pearson r = 0.80 and 0.73, respectively, p-values < 0.05), and revealed that the percentage of high IAV-transcribers among infected cells was stable across populations (**Fig. 5*E***). Interestingly, variation in high/low transcribers had the strongest impact on viral mRNAs in EUB individuals (Pearson r = 0.66 and 0.09, for EUB and AFB, respectively, interaction p-value = 9.5×10^-8^; ***SI Appendix*, Fig. S8*B***). This could be explained by the overall higher rate of infection that we observed in EUB relative to AFB (+9.6 % of infected cells, Wilcoxon p-value= 5.2×10^-10^), together with higher cell death (+5% of dying cells, Wilcoxon p-value = 2.6×10^-10^). Taken together, these results show that while variation in the proportion of high IAV-transcribing infected cells has a sizeable impact on viral mRNA levels, population differences in viral mRNA levels are primarily driven by the overall proportion of cells that will ultimately become infected.

### Supplementary Figures

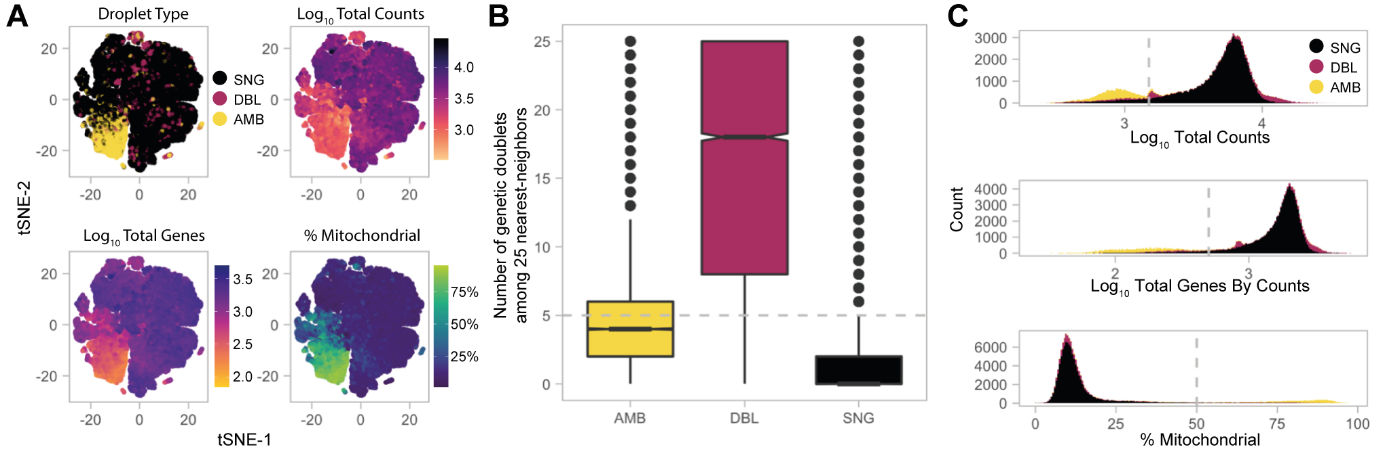

#### Fig. S1. Initial Quality Control of Droplet-Based scRNA-seq Data Excludes Doublets and Low-Quality Cells

(***A***) t-distributed stochastic neighbor embeddings (tSNEs) of 132,130 cell-containing droplets colored by various QC metrics. (***B***) Doublet detection based on genetics and nearest-neighbors. Cell-containing droplets were traced back to donors using two independent methods - Demuxlet and SoupOrCell - both of which capitalize on genetic variation in the sequencing reads. Comparison of genetically deemed doublets to high-confidence singlets (concordant identification of the donor across both programs) revealed that doublets are more likely to share nearest-neighbors in a *knn*-graph with other doublets, and we used this feature to identify droplets presumed to contain two or more cells originating from the same donor. Barcodes with > 5 genetic doublets as nearest-neighbors were considered doublets and excluded from post-QC analyses. (***C***) Distribution of standard QC metrics across all 132,130 barcodes, colored by droplet type. Droplets with < 1500 total counts (top), < 500 genes (middle), or > 50% mitochondrial gene content (bottom) were excluded from further analysis and these thresholds are designated with grey dotted lines. Abbreviations: SNG, singlets; DBL, doublets; AMB, ambiguous

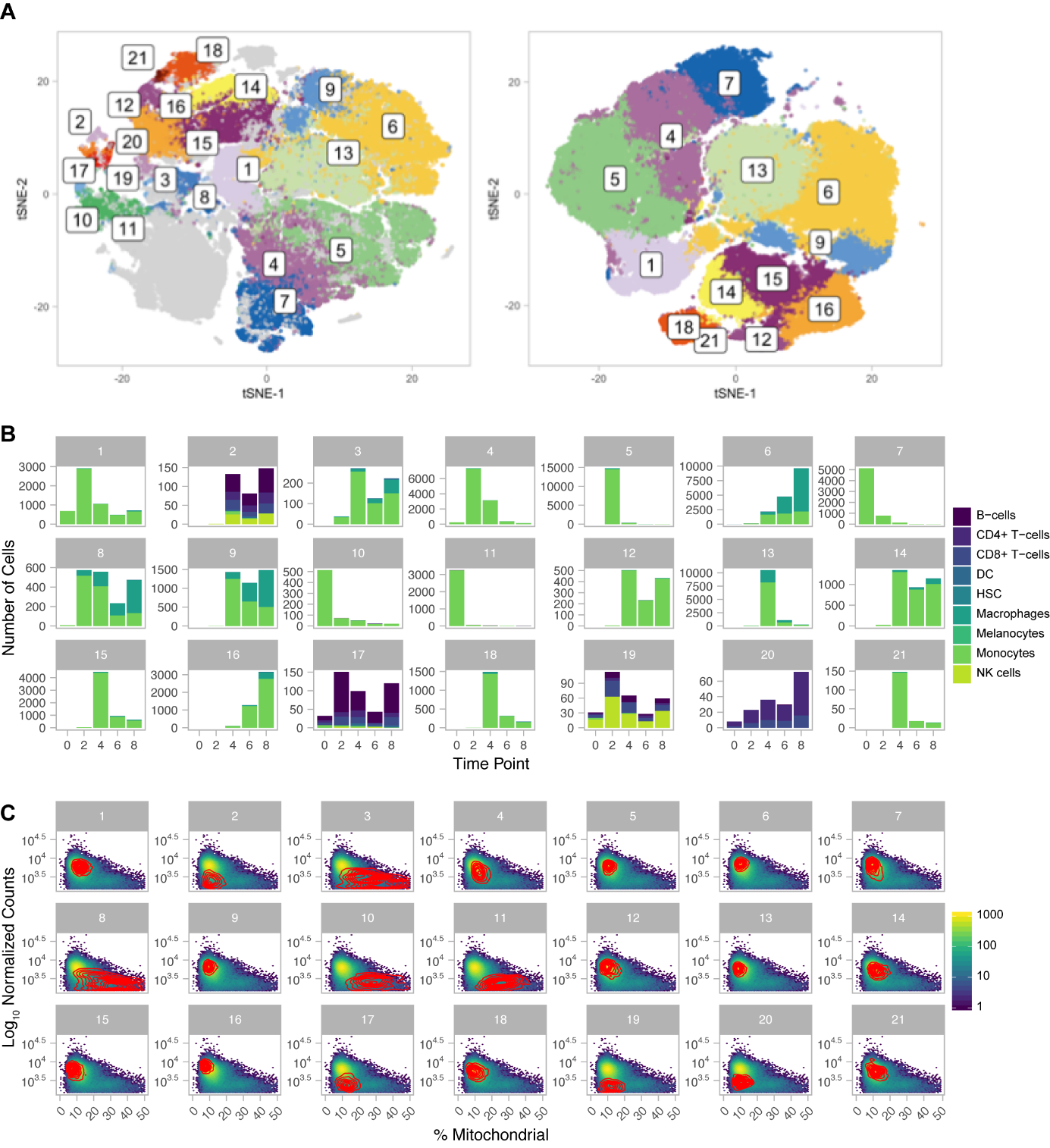

#### Fig. S2. Additional Quality Control of Droplet-Based scRNA-seq Data Excludes Contaminants and High-Quality Dying Cells

(***A***) Pre-QC tSNE (left) and post-QC tSNE (right) colored by graph-based clusters. Cells that were excluded in the first stages of QC (e.g. doublets, of cells with low read counts and high mitochondrial content) are colored in grey on the pre-QC tSNE. (***B***) SingleR cell type predictions for 96,386 high-quality single cells stratified by cluster membership and time point are displayed. We excluded clusters 2, 17, 19 and 20 from further analyses due to the predominance of contaminant cell types. (***C***) Identification of dying cell clusters. 2D kernel density estimation (red contour) is plotted for each cluster against the background of the total dataset represented by the hexagonal heatmap showing the distribution for two quality control metrics for 96,386 high-quality single cells. We exclude clusters 3, 8, 10, and 11. Clusters 2, 17, 19, and 20 are also outliers, but were already deduced to be contaminant cell populations.

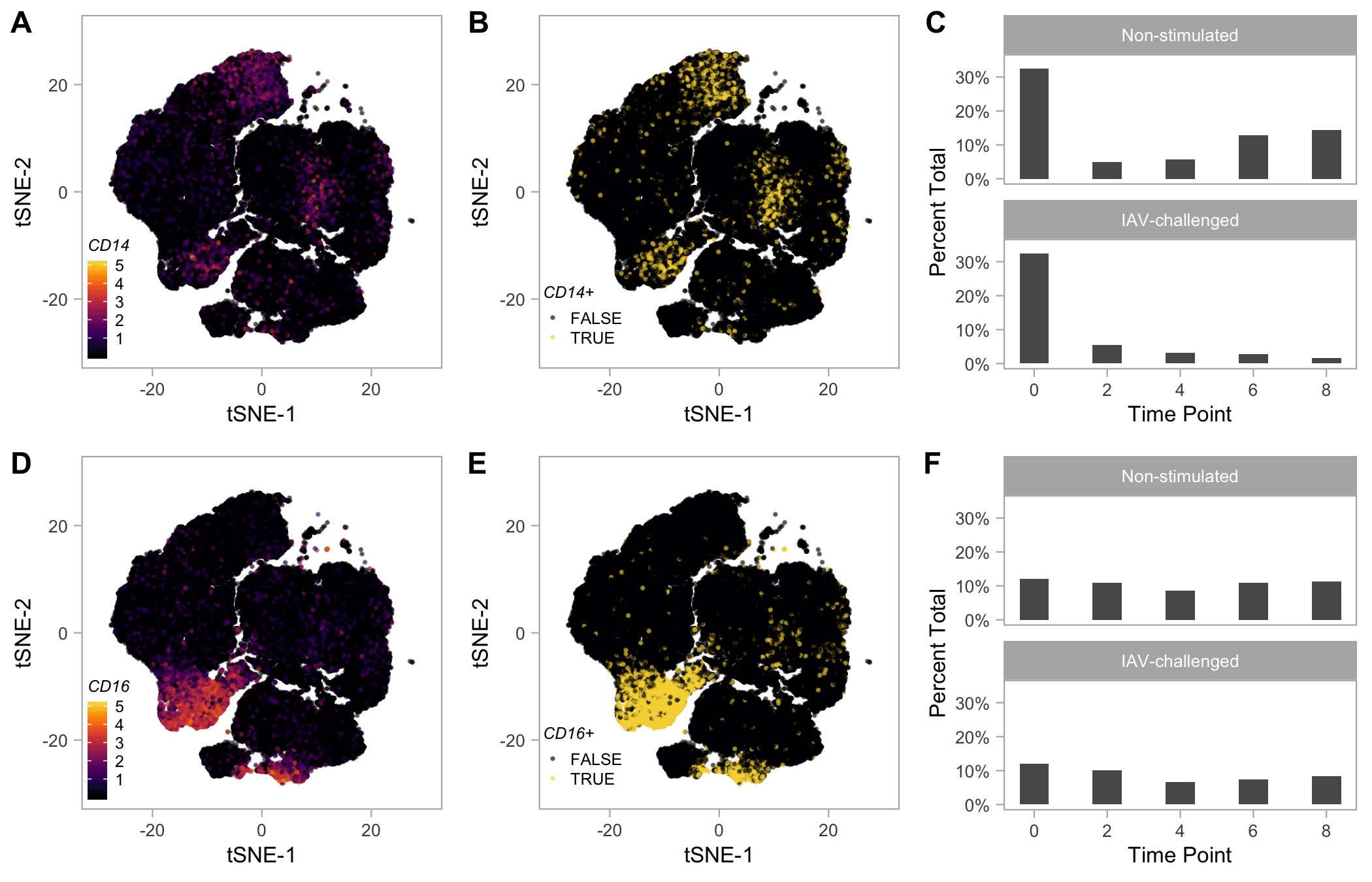

#### Fig. S3. mRNA Expression of CD14 and CD16 (FCGR3A) Highlight Distinct Cell Populations

(***A***) tSNE plot colored by log_2_ normalized counts for *CD14*. (***B***) tSNE plot colored based on significance of *CD14* expression when accounting for ambient RNA (*CD14^+^*, FDR < 0.01). (***C***) Bar plot showing the percentage of *CD14^+^* cells across time points and conditions. (***D***) tSNE plot colored by log_2_ normalized counts for *FCGR3A* (aka *CD16*). (***E***) tSNE plot colored based on significance of *FCGR3A* expression when accounting for ambient RNA (*CD16^+^*, FDR < 0.01). (***F***) Bar plot showing the percentage of *CD16^+^* cells across time points and conditions.

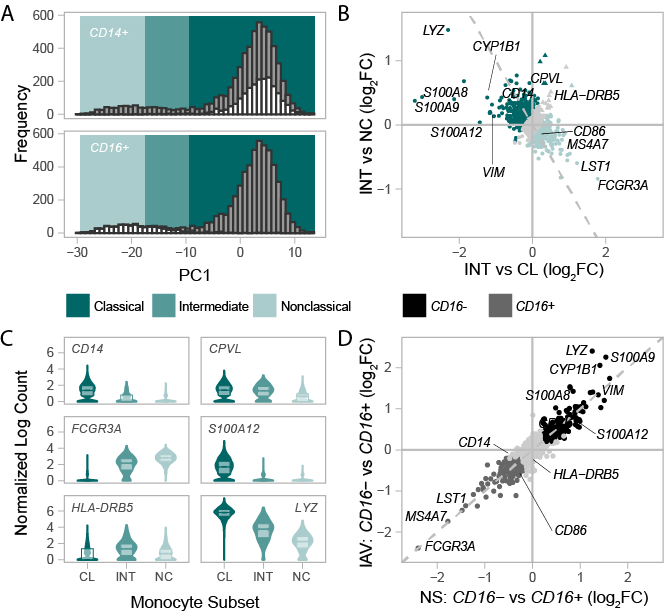

#### Fig. S4. scRNA-seq Accurately Recovers Canonical Monocyte Subsets at T_0_

(***A***) Distribution of 6,601 monocytes along PC1 at T_0_, highlighting *CD14^+^* (top panel) and *CD16^+^* (bottom panel) cells. Differing shades of green distinguish nonclassical (light green), intermediate (medium green), and classical (dark green) monocyte subsets. (***B***) Average gene expression difference among the three canonical monocyte subsets for 4,589 genes at T_0_. The average log_2_FC change in expression between intermediate and classical subsets are plotted on the x-axis, while the average log_2_FC between intermediate and nonclassical subsets are plotted on the y-axis. Triangles designate the 18 genes upregulated in the intermediate subset relative to both classical and nonclassical subsets (log_2_FC>0.2, FDR≤1%). (***C***) Example distributions of gene expression for monocyte subset markers at T_0_. Abbreviations: classical (CL), intermediate (INT), nonclassical (NC), non-stimulated (NS), and IAV-challenged (IAV).

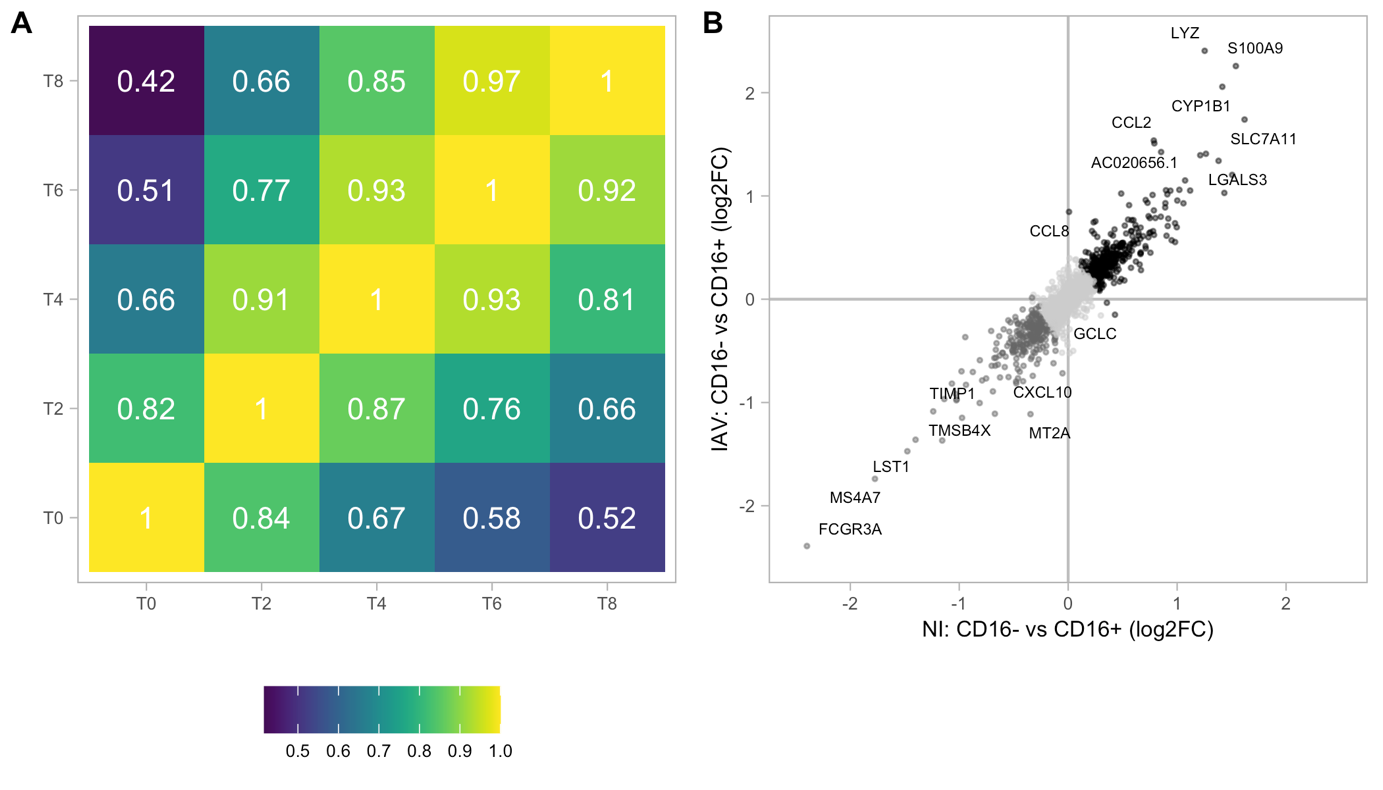

#### Fig. S5. scRNA-seq Accurately Recovers *CD16^+/-^* Subsets and their Functional Roles

(***A***) Correlation among log_2_ fold change in gene expression between *CD16^+/-^* subsets across time points. Upper-diagonal reflects the non-infected condition, while lower-diagonal reflects the IAV-challenged condition. Fill color and text report the Pearson r. (***B***) Average gene expression differences between *CD16^-^* and *CD16^+^* monocyte subsets for 5,681 genes across T_2_ to T_8_ in two conditions. Differences between the two subsets in the non-stimulated condition are plotted on the x-axis, while differences between the two subsets in the IAV-challenged condition are plotted on the y-axis. Genes consistently differentially expressed between *CD16^+^* and *CD16^-^* cells across all time points (including T_0_), conditions, and donors (log_2_FC>0.2, FDR<1%) are highlighted in dark grey (upregulated in *CD16^+^* subsets) or black (upregulated in *CD16^-^* subsets).

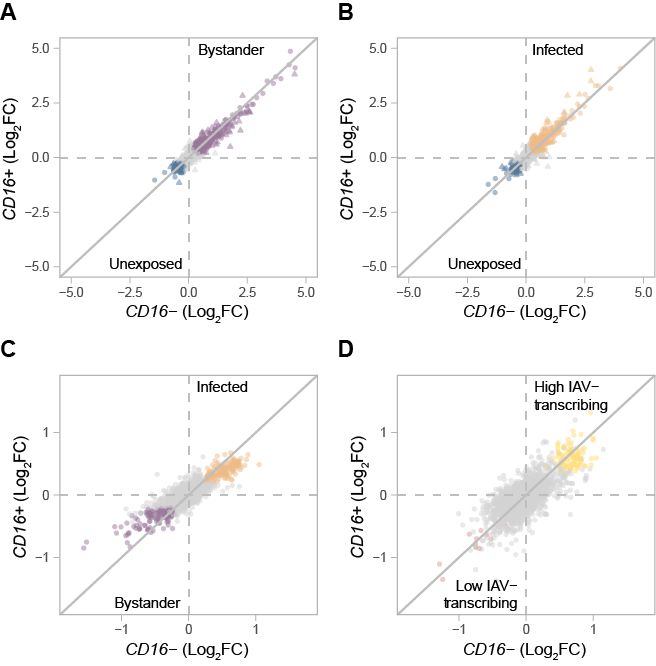

#### Fig. S6. *CD16^+/-^* Subsets Display Similar Transcriptional Responses to IAV

For each cell state comparison, the average log_2_FC in expression between (***A***) unexposed and bystander, (***B***) unexposed and infected, (***C***) bystander and infected, and (***D***) low and high IAV-transcribing infected cells is plotted for *CD16^-^* cells on the x-axis and *CD16^+^* cells on the y-axis. Colors reflect common transcriptional responses among cell states (>0.2 log_2_FC in both subsets, FDR≤1%). Triangles in *A* & *B* represent genes which demonstrate a stronger response in a particular monocyte subset (interaction test, FDR≤1%).

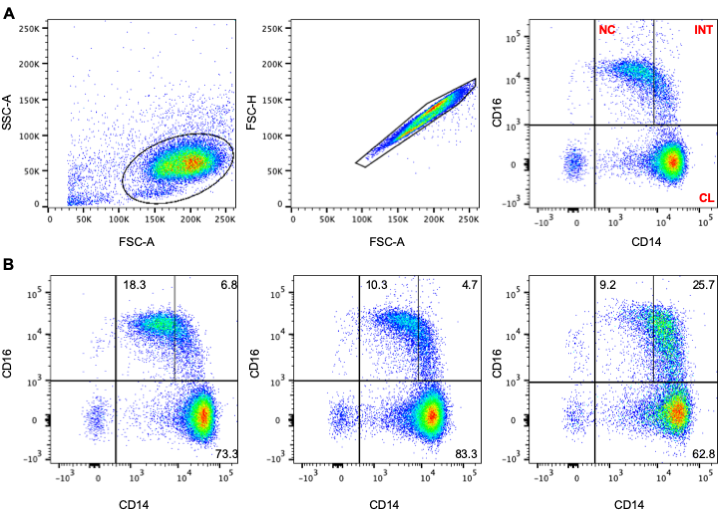

#### Fig. S7. Flow Cytometry Analysis and Gating Strategy of EVOIMMUNOPOP Samples.

(***A***) Gating strategy to discriminate distinct monocyte subsets based on CD14 and CD16 expression, after doublet exclusion with forward scatter-height and forward scatter-area. (***B***) Sample flow cytometry profiles for three donors of the EVOIMMUNOPOP cohort with varying proportions of monocyte subsets. Abbreviations: classical (CL), intermediate (INT), and nonclassical (NC).

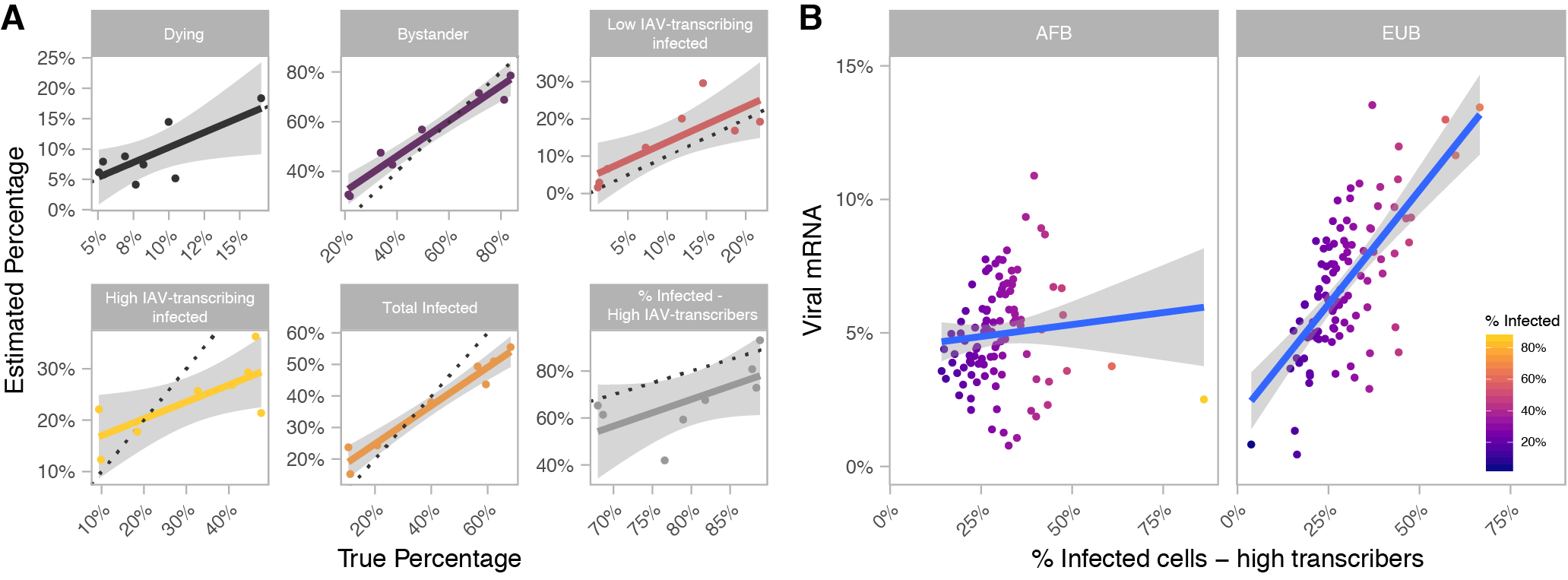

#### Fig. S8. Deconvolution of Bulk RNA-Seq Profiles Correctly Infers Proportion of Infected Cells and Highlights Greater Infectivity among European-Ancestry Individuals

(***A***) Cross-validation of the deconvolution method. For each cell state, comparison between the percentage of UMIs that originate from that cell state in the scRNA-seq data and the percentages estimated from their bulk RNA-seq profiles are plotted. A comparison is also provided for the total percentage of infected cells (orange) and the percentage of high IAV-transcribers among infected cells (grey). For each comparison, a regression line is depicted with a 95% confidence interval. Dotted lines indicate when estimates are identical to the observed values. (***B***) Percentage of IAV reads as a function of the estimated percentage of high IAV transcribers among infected cells, for African and European ancestry individuals (AFB and EUB, respectively). Colors reflect the estimated percentage of infected cells for the sample. One individual with no infected cells was excluded (*n*_AFB_=99, *n*_EUB_=99).

### Supplementary Tables

#### Table S1. Experimental Design for Single-Cell RNA-sequencing Libraries

In each library, samples from both conditions were pooled prior to being run on the 10X Chromium. Donors are labeled D1-D8 based on the rank of their pseudo-bulk viral mRNA content at T_4_. Abbreviations: AFB, African-ancestry individual from Belgium; EUB, European-ancestry individual from Belgium; IAV, challenged with A/USSR/90/1977(H1N1) at a MOI=1; NI, non-infected (control).

| **10X Library** | **Donor** | **Population** | **Condition** | **Time Point** |
| --- | --- | --- | --- | --- |
| L1 | D5 | EUB | NI | 0 |
| L1 | D2 | EUB | NI | 0 |
| L1 | D7 | EUB | NI | 0 |
| L1 | D1 | EUB | NI | 0 |
| L1 | D4 | AFB | NI | 0 |
| L1 | D6 | AFB | NI | 0 |
| L1 | D8 | AFB | NI | 0 |
| L1 | D3 | AFB | NI | 0 |
| L2 | D1 | EUB | IAV | 2 |
| L2 | D8 | AFB | IAV | 2 |
| L2 | D3 | AFB | IAV | 2 |
| L2 | D5 | EUB | NI | 2 |
| L2 | D7 | EUB | NI | 2 |
| L2 | D6 | AFB | NI | 2 |
| L3 | D5 | EUB | IAV | 2 |
| L3 | D7 | EUB | IAV | 2 |
| L3 | D6 | AFB | IAV | 2 |
| L3 | D2 | EUB | NI | 2 |
| L3 | D4 | AFB | NI | 2 |
| L3 | D3 | AFB | NI | 2 |
| L4 | D2 | EUB | IAV | 2 |
| L4 | D4 | AFB | IAV | 2 |
| L4 | D6 | AFB | IAV | 2 |
| L4 | D5 | EUB | NI | 2 |
| L4 | D1 | EUB | NI | 2 |
| L4 | D8 | AFB | NI | 2 |
| L5 | D7 | EUB | IAV | 4 |
| L5 | D1 | EUB | IAV | 4 |
| L5 | D4 | AFB | IAV | 4 |
| L5 | D6 | AFB | NI | 4 |
| L5 | D8 | AFB | NI | 4 |
| L5 | D3 | AFB | NI | 4 |
| L6 | D5 | EUB | IAV | 4 |
| L6 | D2 | EUB | IAV | 4 |
| L6 | D6 | AFB | IAV | 4 |
| L6 | D7 | EUB | NI | 4 |
| L6 | D1 | EUB | NI | 4 |
| L6 | D4 | AFB | NI | 4 |
| L7 | D4 | AFB | IAV | 4 |
| L7 | D8 | AFB | IAV | 4 |
| L7 | D3 | AFB | IAV | 4 |
| L7 | D5 | EUB | NI | 4 |
| L7 | D2 | EUB | NI | 4 |
| L7 | D1 | EUB | NI | 4 |
| L10 | D2 | EUB | IAV | 6 |
| L10 | D1 | EUB | IAV | 6 |
| L10 | D6 | AFB | IAV | 6 |
| L10 | D4 | AFB | NI | 6 |
| L10 | D8 | AFB | NI | 6 |
| L10 | D3 | AFB | NI | 6 |
| L8 | D7 | EUB | IAV | 6 |
| L8 | D4 | AFB | IAV | 6 |
| L8 | D3 | AFB | IAV | 6 |
| L8 | D5 | EUB | NI | 6 |
| L8 | D2 | EUB | NI | 6 |
| L8 | D8 | AFB | NI | 6 |
| L9 | D5 | EUB | IAV | 6 |
| L9 | D2 | EUB | IAV | 6 |
| L9 | D8 | AFB | IAV | 6 |
| L9 | D7 | EUB | NI | 6 |
| L9 | D1 | EUB | NI | 6 |
| L9 | D6 | AFB | NI | 6 |
| L11 | D7 | EUB | IAV | 8 |
| L11 | D1 | EUB | IAV | 8 |
| L11 | D3 | AFB | IAV | 8 |
| L11 | D5 | EUB | NI | 8 |
| L11 | D2 | EUB | NI | 8 |
| L11 | D4 | AFB | NI | 8 |
| L12 | D5 | EUB | IAV | 8 |
| L12 | D4 | AFB | IAV | 8 |
| L12 | D6 | AFB | IAV | 8 |
| L12 | D7 | EUB | NI | 8 |
| L12 | D8 | AFB | NI | 8 |
| L12 | D3 | AFB | NI | 8 |
| L13 | D2 | EUB | IAV | 8 |
| L13 | D8 | AFB | IAV | 8 |
| L13 | D3 | AFB | IAV | 8 |
| L13 | D7 | EUB | NI | 8 |
| L13 | D1 | EUB | NI | 8 |
| L13 | D6 | AFB | NI | 8 |
